# Tactical signalling by victims increases bystander consolation in bonobos

**DOI:** 10.1101/2022.01.18.476740

**Authors:** Raphaela Heesen, Diane A. Austry, Zoe Upton, Zanna Clay

## Abstract

Tactical emotion communication has long been considered uniquely-human. As a species, we readily exaggerate, inhibit and modify emotional expressions according to social context and audience. Notably, emitting emotional displays, such as those pertaining to distress states, can evoke empathic responses in others such as the offering of consolation to victims after a fight. Animal emotion expressions, by contrast, are traditionally viewed as uncontrollable arousal responses. Our study challenges this view by assessing the level of control in the emotional signalling of sanctuary-living bonobo victims following aggressive attacks (N = 27 victims, *N* = 144 attacks) and its and its corresponding effect on receivers. Results show that the production of paedomorphic signals by adult bonobo victims increased chances of receiving consolation from bystanders and reduced risk of future aggression from former opponents, highlighting a strategic function. Victim signalling also increased with audience size, yet strategies differed by age: immature bonobos were more likely to cease signalling in proximity of close-social partners, whereas adults were more likely to cease signalling after having been consoled. These data suggest that bonobo emotion communication has a developmental trajectory and that tactical emotion signalling is a *Pan-human* capacity, preceding the split of *Homo*.

## Introduction

Compared to other species, humans exhibit impressive levels of control in our expressions of emotional states. This apparent cognitive control over emotion signalling has been suggested to play a crucial role in the emergence of modern human societies and language, and more generally underpins our social interactions and how we relate to one another [1, 2]. As a species, we not only regularly modify and exaggerate emotional expressions depending on our social environment, but we also inhibit or cover our emotional states to influence others and prevent others from exploiting us. Emotional control therefore allows us to maintain a complex social life by enabling us to flexibly adapt to social situations and audiences, achieve social goals, and successfully collaborate [3, 4]. Crucially, effectively expressing emotions and desires triggers emotional responses in receivers and in this respect, affective signalling has been considered as a form manipulation or influence over receiver responses [5]. For instance, signalling distress can result in the bystanders witnessing the event offering empathic comforting to the signaller, which in turn, can alleviate stress and further strengthen social bonds [6, 7].

While humans seem particularly adept at tactically adjusting their emotional signalling behaviour, relatively little is currently understood about how humans have evolved such impressive forms of communicative control, and the extent to which such control is a derived feature of our species or more evolutionarily ancient [e.g., 2].

Comparative data, including from our closest living great ape relatives, are crucial to gather insights into the evolution of complex emotion signalling of our own species, and broaden our perspective about emotional intelligence of nonhuman animals (henceforth animals) more generally.

In humans, the expression of distress through crying represents a good example for emotion communication that presents both an involuntary and voluntary dimension. Although the perception of distress is better understood than its expression [8,but see 9], it is evident that crying is a complex emotion expression that develops increasing cognitive control and awareness across the lifespan. Expressions of negative affect, like human crying, are particularly strong drivers of social attention in receivers; moreover, such expressions are likely to directly impact on receiver responses, including the prosocial offering of consolation, which refers to friendly contact that functions to alleviate the distress in the recipient [10]. Research with young human children suggests that attention towards distress signals is especially pronounced when the distress occurs as a result of evident harm (as compared to distress for no obvious reason), or if the harm has been committed by the receivers themselves [11]. Although crying before the age of 7-9 months in human infants is thought to primarily function to promote caregiver proximity and caregiving to increase infant survival, crying increasingly involves greater cognitive awareness, that [expanding especially with language 12]. This is also evident in the later expansion of its functionality, by which human infant crying can promote peer support, for example in the form of reconciliation and consolatory behaviours [13, 14]. At the preschool age, children become more proficient at regulating crying, along with other negative and positive emotion expressions, as well as using language to explicitly inform others about their feelings [8,12,15].

Emotion expressions in humans, like crying, are naturally expressed and recognized via cross-modal integration of facial expressions, vocal utterances and gestures, where gestures seem more easily controllable and might be used to add and disambiguate meaning [16, 17]. Intriguingly, such cross-modal integration within the context of emotional exchange has also been demonstrated in the vocal-gestural signal combinations of our close relatives, the bonobos [18, 19], suggesting such capacities are not human unique. Nevertheless, the evolutionary basis of emotional control is relatively unexplored in the animal research literature.

Until recently, the notion of even studying emotions, and related constructs such as empathy, in animals has been somewhat taboo, and whether or not animals can control their emotion expressions remains controversial [20, 21]. Traditionally, animal communication has been viewed as “emotional-by-default”, with *emotional* denoting uncontrollable, involuntary, arousal-based and overall, scientifically inaccessible.

According to this view, animals predominantly react to certain events by expressing arousal states and lack intentional control or awareness about their emotional state and its potential effect on receivers (e.g., an alarm call given towards a predator). The signaller is also generally assumed to stop signalling after the stimulus has ceased, regardless of whether the audience has or has not perceived the expression. By contrast, human emotional communication can be both - emotional *and* intentional [see for discussions 1,20,22,23].

With a call to the apparent rise of an era of ‘affectivism’ in human sciences, from the long-standing influence of behaviourism and cognitivism [24], scholars are increasingly criticizing the dichotomous view of emotion versus cognitive communication, including in animals [1, 20]. This includes stating that *i)* emotion communication may be both affective and cognitive simultaneously, and that *ii)* animals, notably nonhuman primates, can flexibly adjust their signals, even when these signals are emitted in emotional settings [1]. For example, great apes can intentionally control their communication as a means to reach social goals [25] or to deceive others [26]. Especially in urgent contexts like contest, predation or aggression, great apes can increase or modify the production of their vocalisations, facial expressions, gestures and body signals depending on the audience members [27], and in some cases to inform [28] or to manipulate recipients’ behaviour and attentive state [19, 29]. Apes are also known to offer empathic comfort to peers expressing distress, such as to victims following social conflicts[10, 30]. While the decision to offer comfort to distressed victims is presumably influenced by the nature of the victim signals themselves, the extent to which such distress signals may be produced strategically to influence bystander prosocial responding remains unknown.

While intentionality appears to be a more general feature of great ape communication, the fact that gestures and body signals are by default operationalised as intentional signals [e.g., 31] whereas vocalisations and facial expressions [e.g., 32] are not creates a bias in the literature, leading to the false assumption that only gestural behaviours can be controlled [1, 33]. Nevertheless, regardless of modality, the level of cognitive control involved in the communication of emotions in particular, as well as the function of controlled emotional signalling, remains unknown. Furthermore, although such signals may facilitate prosocial responses by receivers (such as offering consolation to victims, see Photo Panel 1), comparative evidence of this remains obscure. Such data would be crucial, however, to inform on the evolution of emotional responding, particularly empathy, and to identify potential selective forces shaping emotional capacities in the Pan-human lineage.

New evidence focusing on the role of cognition in great ape emotion signalling [28, 29] warrants investigation into the degree of intentionality in emotional communication. For a number of reasons, bonobos (*Pan paniscus*) [see for review 34], represent a particularly promising primate model both to assess the degree of control in signaller emotion expressions, as well as the extent to which such signals shape the empathic responses of bystanders [35]. This is due to their apparently heightened levels of social tolerance [36], strong social orientation and sensitivity towards socio-emotional cues [37–39], increased awareness about social partners and commitments [40, 41] and prosociality [42], even towards outgroup individuals [43]. Bonobos are also known for their paedomorphic traits, such as playfulness even in adulthood [44, 45] and morphological features like smaller canine teeth and juvenilized cranium [46], which have been suggested to result from an evolutionary process of self-domestication [46].

Importantly, bonobos have been documented to exhibit impressive levels of empathy towards distressed conspecifics as demonstrated by their tendency to approach victims in distress to offer them comforting contact [47, 48].

The first aim of the current study is therefore to assess the level of control involved in emotion signalling of bonobos during a high-arousal context, where volitional control may otherwise be presumed to be low. The second related aim is to investigate the function of such emotion signals with respect to the victim’s potential goals. To address these aims, we examined the communicative signalling of bonobo victims after naturally occurring social conflicts; we tested the extent to which different forms of victim signalling predicted bystander prosocial responses [e.g., 10]. In doing so, our broader goal is to address the evolutionary origins of emotion control in our own species, as well as to assess the validity of the theory that emotion control precedes the emergence of human sociality and language [1, 2]. Given that emotional signalling in humans can be controlled and has a strategic function, including in the elicitation of prosocial empathic responses in receivers, we hypothesized that if bonobos are somewhat cognizant of their own emotion signalling, they should adjust their signalling patterns according to the nature of their audience and likelihood of receiving assistance.

As well contributing novel insights into the nature of emotional communication in primates, such findings of strategic victim signalling would also challenge the current assumption that consolation, and other expressions of empathy in animals, are bystander-initiated behaviours, rather than being elicited by the signaller themselves [49]. In fact, most comparative studies examining consolation in animals, including primates, specifically exclude victim-initiated bystander affiliation, because such affiliation is thought to be driven by the victim themselves rather than representing genuine empathic motivation by the bystander [35, 49]. Addressing victim signalling and its potential role in shaping the affiliative responses of bystanders (consolation) and former opponents (reconciliation) therefore addresses a crucial ignored component of post-conflict interactions, and will inform on whether consolation in bonobos is truly spontaneous or instead may be mediated by tactical emotion communication of the victims themselves.

If bonobo victims’ distress expressions are more than indicators of negative arousal with little or no underlying cognitive control, we would expect that bonobos deploy certain signalling techniques to achieve different social goals. These goals might be related to, for instance, receiving *consolation* from bystanders (e.g., see photo 1), repair of relationships with former opponents via *reconciliation*, or *prevention of future attacks* from former opponents and other group members. To allow for a multi-component and multi-modal analysis [50] of victim signalling, we here assessed the use of vocalisations, facial expressions, gestures (manual movements produced with the limbs and head) and body signals (movements of the entire body) of victims.

**Photo Panel 1.**
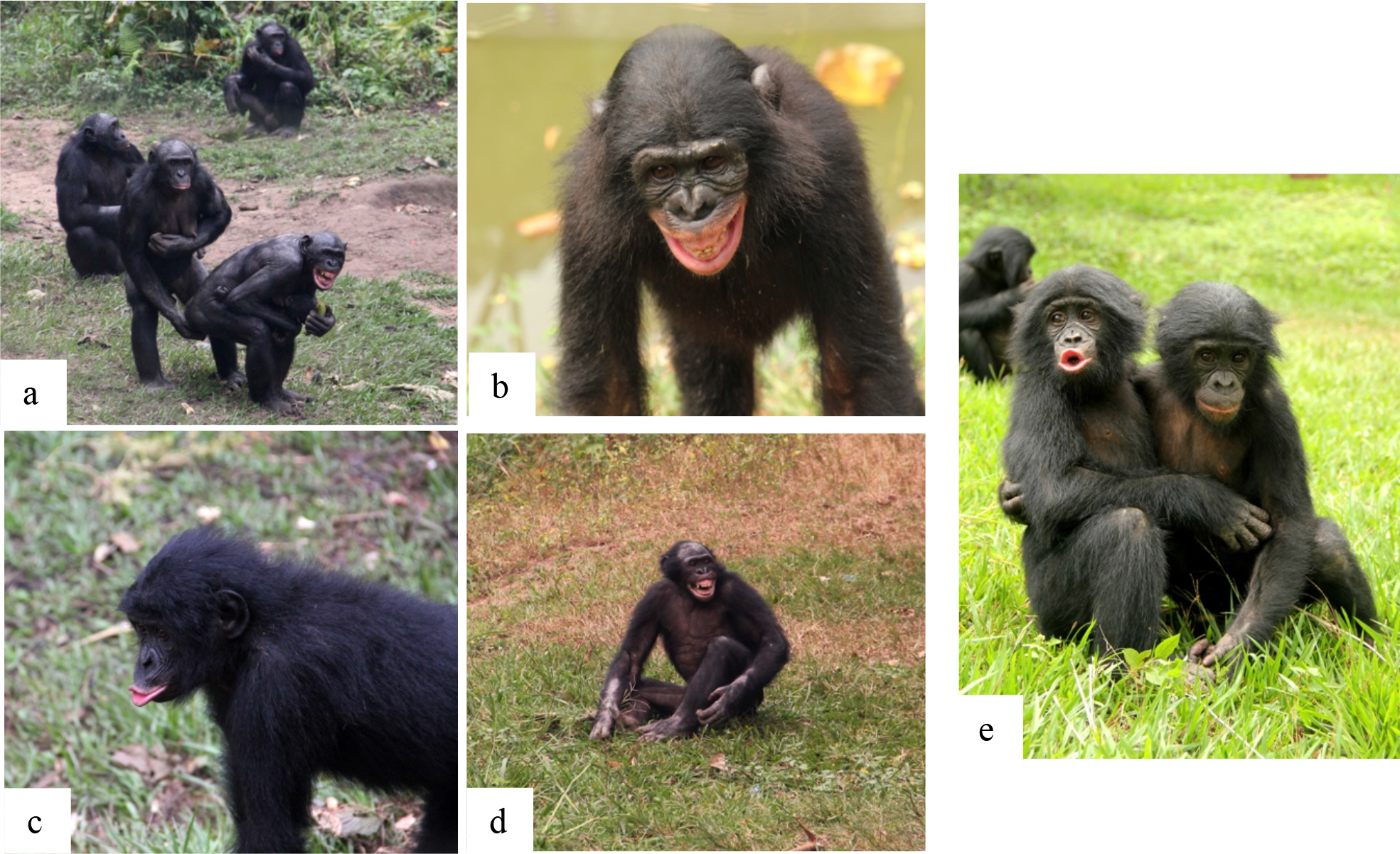
Photographs depicting emotion expressions of bonobo victims following social conflicts, taken at Lola ya Bonobo Sanctuary. a) Adult female victim crouching with bared teeth face expression/scream vocalisation, being consoled by an adult female; b). Example of bared-teeth facial expression, c) example of pout-face expression, d) example of victim scream/bared teeth plus piloerection; d) example of victim with pout face being consoled by a juvenile bystander © Zanna Clay/ Lola ya Bonobo Sanctuary.

To further discriminate between functions of different signalling styles, we distinguished between three principal signal categories: (1) *paedomorphic signals;* i.e. those resembling the signals typically used by immature bonobos [18], *aggressive signals,* those used during tense situations or conflicts [19] and general *affiliative-submissive* signals, those signals thought to signal submission or appeasement of dominants [51]. In humans, paedomorphic appearance and behaviours, in particular, are connected with innate mechanisms that stimulate caregiver protection and assistance [52–54].

Therefore, we predicted that the occurrence of paedomorphic signals (aka “juvenilised” signals, frequently seen in infants) by bonobo victims should positively predict the occurrence of prosocial responses by bystanders - via consolation - and by former opponents, via reconciliation, as compared to other signals. Paedomorphic signals should also be expected to reduce the risk of renewed aggression (prediction A- paedomorphic signals). Likewise, since aggressive behaviours (e.g., physical slapping, hitting, kicking, aggressive barks/screams) are generally more likely to repel future aggressors [27], we expected that aggressive signals by bonobo victims following conflicts should be associated with reduced risk of renewed aggression (prediction B- aggressive signals).

Moreover, if emotional signalling functions as a tool to promote consolation from bystanders and/or reconciliation by former opponents, we predicted that bonobos should cease signalling after having received either of these post-conflict affiliative contacts (prediction C- goal sensitive signalling).

Additionally, if bonobos take into account their audience while signalling [55], we should expect bonobo victim signalling to vary according to audience size and composition, regardless of the severity of the attack or the victims’ arousal level. Here, we assessed victim arousal by denoting the occurrence of piloerection of the fur, considered to be a valid indicator of arousal in primates e.g.,55]. If bonobo victims are capable of tactically tailoring their emotion signalling to the size and composition of the audience, we expect that victims should increase the complexity and saliency of victim signalling, as measured by number of signal components (gestures, vocalisations, facial expressions, body signals) and overall signalling display duration, in the presence of larger audiences, particularly those containing close-social partners (prediction D- audience size and composition). Indeed, previous research on the likelihood of chimpanzees’ signalling persistence showed that persistence increases as a function of audience size [57].

Alternatively, if close-social partners (henceforth friends) are already more likely to console victims, regardless of signalling strategy, we would expect victims should decrease signalling efforts when friends are present who can offer support (prediction E, see [57]). In other words, friends or “allies” are assumed to provide more support than non-allies [47] which should be less contingent on signalling effort.

Since in humans, communication about and responses to distress increase in complexity with age, we also investigated the developmental trajectory of victim signalling in our study sample. In particular, given limitations in cognitive maturation, we expected immature bonobos would be less proficient at adapting their signalling based on audience and social goals. Specifically, we expected that compared to adults, immatures would show less evidence of adjusting their signalling techniques to enhance chances of consolation, reconciliation or reduce renewed aggression, be less likely to stop signalling even when consolation and reconciliation were already provided, and be less likely to adjust signalling efforts to audience size and composition.

## Methods

### Study subjects and site

Observations of naturally-occurring victim signalling behaviour were conducted on two groups of bonobos living at the Lola ya Bonobo Sanctuary, in Kinshasa (Democratic Republic of Congo), during two study periods. In May-August 2011, N = 301 observation hours were collected at group 1 comprising 25 individuals; and N = 152 observation hours group 2 comprising 17 individuals. From May-August 2012, N = 205 observation hours of group 1 comprising 22 individuals and N = 187 observation hours of group 2 comprising 20 individuals were conducted. For details of group composition in 2011-2012 see [47, 48]. Most bonobos living in the sanctuary were orphaned victims rescued from the bush meat trade who had been reared by a human mother and then been reintegrated in a social group of conspecifics at the sanctuary. Individuals born at the sanctuary remain with their mothers in their social group. The bonobos ranged among three large, forested enclosures (15-20ha) throughout the day, and voluntarily slept inside dormitory shelters (approx. 75m^2^, split into several open sub-rooms) during the night. The bonobos received food 3-4 times a day from their respective caregivers, with a variety of fruits and vegetables.

### Data collection

Naturalistic observations were recorded during the day, when bonobos were ranging outside in their forested enclosures. We analysed a total of N = 150 conflicts, of which N=6 were excluded either due to insufficient visibility for analysis of victim signals or due to human-related conflicts. Conflicts involved N = 27 victims [females: 12; males: 15; immatures < 10 years =14; adults ≥ 10 years =13] and N = 23 aggressors [females: 11; males: 12; immatures (< 10 y) = 6; adults (≥ 10 y) =17] from across the two groups, see Table S1 A & B. We denoted adults as those individuals becoming (or being) sexual mature above the age of 10 years; immatures were individuals being infant, juvenile, or young sub-adult below the age of 10 years (Table S1).

**Table 1.** Overview of communication styles.

| Communication style code | Full name | Characteristics of signal types within this classification |
| --- | --- | --- |
| **P** | Paedomorphic signals | Signals with juvenile features, often seen in immatures of young age in play or requests-to-mother, such as tantrum behaviours, pout moan vocalisation, pout face and reach gestures. |
| **AF** | Affiliative submissive signals | Signals making victims appear submissive, appease the opponent, having an inhibiting effect on the partners' aggressive tendencies, or which contain affiliative components, such as soft and tactile gestures, crouching, bared-teeth expression and victim screaming. |
| **AG** | Aggressive signals | Harsh signals to threaten or indicate counter attack towards opponent or other group members (aggression redirection); e.g., rough gestures / threat barks / chase-galloping attempts. |

We conducted all-occurrence observations of agonistic interactions that included at least one of the following behavioural elements [methods detailed in 48]: recipient fleeing and/or screaming in reaction to aggression, aggressor threat barking/grunting, directed display charge, threat arm wave, chase, hit, trample, slap, shove, poke, or bite. For each conflict, we recorded the identities of the victim and the aggressors, as well as the identities of all visible bystanders. We also recorded whether bystanders were physically present within 5 m, 5–10 m or beyond 10 m of the victim at conflict start.

Following [47, 48], we video-recorded (and later analysed) post-conflict periods of 5 minutes duration using a Canon Vixia HF200 HD camcorder with a Sennheiser microphone (MKE400) attached (i.e., any behaviour after 5 min was not included in the analysis). To assess social relationships, we also conducted instantaneous scan samples at 10-min fixed intervals throughout the day, in which we recorded the identities of all individuals engaged in any affiliative behaviours (and of respective interaction partners) such as grooming, contact sitting, sitting within arm-reach proximity, playing, or engaging in sexual contact.

### Video coding

Videos were coded in ELAN (vs. 5.7) [59]. Each conflict was considered individually. For each conflict, we coded for study year, post-conflict (PC) ID, victim identity/aggressor identity, study group, aggression severity, PC events (consolation, reconciliation, renewed aggression), whether victims were piloerect (i.e., fur on the neck or back visibly bristling, see Supplementary Image S1), victims’ signalling display duration, the number of signals, their vocalisation/gesture/body signal/facial expression type and class, and their signalling persistence (whether victims continued signalling in further bouts depending on whether having or having not received consolation/reconciliation).

For the signalling persistence analysis (see specifications in section “statistical analysis”), we coded reception of consolation within after 1 min after the beginning of the bout (or until the next bout if another bout followed within 1 min) and reconciliation within 1 min after the beginning of the bout (or until the next bout if another bout followed within 1 min). If two post-conflict events followed the bout within 1 min after its start, only the first event type was noted, as this was interpreted to have an immediate effect on the signaller’s behaviour; e.g. if a bout was followed by reconciliation after 5 sec of its start and then by consolation after 56 sec after its start, we would have noted the post-bout event as reconciliation for that particular bout. Examples of how videos were coded and for victim post-conflict signalling can be found in supplementary movies s1-s5 (https://figshare.com/s/7dddfc02c919ec4574ef).

### Aggression severity

We distinguished between mild and severe aggression received by victims following [ 25]. Severe aggression occurred when aggressors physically attacked victims by slapping, kicking, shoving or biting them, or when aggressors chased (pursued) victims for more than 7 metres. Mild aggression occurred when the case pursuit was less than 7 metres and when aggressors displaced victims without physically touching them, for example by shaking, throwing or aggressively moving vegetation, producing postural/gestural threats like attempting to chase, dragging objects or slapping the ground or objects.

Post-conflict (*PC) events.* PC events could contain consolation, reconciliation [47, 48] or renewed aggression [27]. Following Clay & de Waal [47, 48], we denoted consolation as being when uninvolved bystanders of the PC *initiated* affiliative contact by physically approaching the victim to offer them friendly physical contact; this included embracing, sexually engaging with, touching, playing or sitting in bodily contact with the victim. We denoted reconciliation as being when former aggressors initiated by approaching the victim to offer affiliative gestures (e.g., head nodding, hand reaching) or contact, including any behaviours described above for consolation towards the victim; affiliative gestures were counted here because aggressor’s behaviours generally seemed to have a large impact the victim’s successive behaviour. Hence, we accounted for any friendly contact directed from the aggressor to the victim in our analysis. In this respect we excluded cases where the victim physically approached (initiated) any post-conflict contacts. We denoted renewed aggression as being when former aggressors re-attacked victims by displacing, taking away resources, chasing, threatening, or physically attacking the victim.

### Signalling display duration

A signal display (and its duration) comprised all signal bouts following aggression until 2 min after the last signal was given, within a max. 5 min window after the first aggression of the victim. A bout is defined as a signalling attempt by the victim towards the aggressor or bystanders followed by a 5 sec response waiting gap (no further signals following) [60].

### Number of signal components (hereafter “number of signals”)

We counted the number of signals in each conflict by summating all single signal components victims produced within the display duration. This meant that each signal component was counted as one signal, with signal components being either a vocalisation, gesture, body signal, or facial expression. For instance, if the victim produced multicomponent signals including a scream overlapping with a scream face, followed by a hand reach gesture that overlaps partly with an arm raise gesture, followed by a bared-teeth expression and a concave back present, we counted six individual signal components. This gave us an idea about the effort in producing signals with several components while keeping a degree of objectivity over what receivers would consider as one or several signals (which cannot be assessed with our data).

### Signalling persistence after consolation/reconciliation

We coded signalling persistence after consolation and reconciliation if victims produced further signal bouts, even after having received consolation or reconciliation within 1 min after the bout, respectively. If instead victims had ceased signalling after these events we coded this as not persisting. Signalling bouts were defined as attempts of signalling towards bystanders or former aggressors with a response waiting period following that was at least 5 sec [60].

### Signal types

To allow for a comprehensive and inclusive analysis of emotion signalling in bonobos, we considered all possible signal components and types [1], see Photo Panel 1 for examples. We followed contemporary great ape communication literature [19,32,51,61–64] and produced an ethogram (see Table S2) of all gesture, vocalisation, facial expression and body signal types recorded in this study. Gestures and body signals followed at least one intentionality criteria of response waiting, audience checking and signal persistence if the goal was not met [e.g., see 65]; vocalisations and facial expressions were coded as they occurred.

### Communication style

Similar to other studies analysing bonobo signals [19] and to allow for a more informative analysis of victims’ communication styles, we divided signal types (gestures, vocalisations, body signals and facial expressions) into three categories based on their intensity, form and – in the case of gestures and body signals – movement and speed. We distinguished between what we called: paedomorphic (P), affiliative- submissive (AS) and aggressive signals (AG), see definitions in Table 1. The distinction between these classes relied on reports from the literature, in which signals had been reported as frequent and typical in a certain age class [P – signals, e.g. ,51,62,66], as being either gentle or aggressive [AG- signals; e.g. ,19,27], or as being affiliative or submissive [AS- signals, e.g. ,51,67–69]. For a full list of each signal types’ category of communication style and proportion of use across individuals, see Table S2-S3.

Signals with juvenile features, often seen in immatures of young age in play or requests-to-mother, such as tantrum behaviours, pout moan vocalisation, pout face and reach gestures.

Signals making victims appear submissive, appease the opponent, having an inhibiting effect on the partners’ aggressive tendencies, or which contain affiliative components, such as soft and tactile gestures, crouching, bared-teeth expression and victim screaming.

Harsh signals to threaten or indicate counter attack towards opponent or other group members (aggression redirection); <u>e.g., rough gestures / threat barks / chase-galloping attempts.</u>

### Coding reliability

The inter-rater test between the two primary coders revealed good agreement for the number of signal types following ethogram Table S2 (N= 16 out of 144 conflicts, 11% of the dataset, Intraclass correlation coefficient ICC = 0.88, N=17). We also computed inter-rater reliability for coding events of consolation and reconciliation. A third coder coded whether there were (or not) events of consolation or reconciliation after conflicts in 20% of the dataset (29 of 145 conflicts). The inter-rater test between main coder and third coder revealed almost perfect agreement for consolation (kappa = 0.93, SE = 0.1) and reconciliation (kappa = 0.92, SE = 0.1). More information about coding reliability can be found in the Supplementary text 1.

### Dyadic bonds

To assess the strength of the dyadic bonds between victims and any of the potential bystanders, we first calculated affiliation scores from the 10 min affinity scans and then identified friends via the upper quartile of the distribution of victims’ affiliation scores with bystanders. Specifically, affiliation scores were computed by the total number of times victims engaged in affiliative behaviours with a certain bystander (contact sitting, grooming, play, sex), divided by the total number of scan samples in which both these partners were present (hence in which they potentially could have affiliated). We then computed a distribution of these proportions for each victim, and identified victims’ close friends by comparing their proportions of affiliations as per shared scans with the upper quartile of the distribution. For any victim, mothers’ affiliation scores were excluded from the calculations to avoid bias of kin relationships with friendship relations.

### Statistical analysis

We used Bayesian mixed models to test our predictions [70]. We characterized uncertainty by two-sided credible intervals (95% CrI), denoting the range of probable values in which the true value could fall. Evidence for an effect in a certain direction (positive or negative) was thus present if posterior distributions shifted away substantially from zero in one direction, as opposed to centring on zero (i.e., which would imply the null expectation of posterior distributions). For inference, we calculated 95% credible intervals from the posterior distributions and checked whether 0 was included.

Additionally, we state the size of the estimated mean of the posterior distribution [parameter estimate b, 70], to assess the association between model predictors and outcome variables. A positive value for *b* would indicate that the subjects exhibit a *higher* probability to produce a certain behaviour when the predictor variable scale increases. A negative value for *b* would indicate the opposite. We also assessed the standard deviation (SD) of posterior distributions [70] and checked how the posterior predictive distributions fitted the empirical response variables (Fig. S2).

To test predictions A-E, we fitted Bayesian generalized and linear mixed models using the Stan computational framework (http://mc-stan.org/), accessed through the brms package [70] (v 2.9.0) in R v. 3.6.1 [71]. Each model included four Markov chain Monte Carlo (MCMC) chains, with 10,000 iterations per chain, of which we specified 2,000 iterations as warm-up to ensure sampling calibration. The model diagnostics revealed an accurate reflection of the original response values by the posterior distributions, since R-hat statistics were <1.05, the numbers of effective samples >100, and MCMC chains had no divergent transitions (see Table S4; Fig S1). We used default priors (weakly informative with a student’s *t*-distribution of 3 degrees of freedom and a scale parameter of 10), see Table S4.

### Predictions A-B: Models 1.1, 1.2 and 1.3

We fitted three generalized mixed models to analyse whether paedomorphic signals increased the likelihood of consolation and reconciliation, and reduced risk of renewed aggression (A), and whether aggressive signals reduced the risk of renewed aggression (B). The dependent variables were binary outcomes of consolation (model 1.1), reconciliation (model 1.2) and renewed aggression (model 1.3) within 5 min post-conflict, fitted with a Bernoulli distribution. The predictors in all models were aggression severity (mild/ severe), piloerection (no/yes), use of paedomorphic signal types (no/yes), use of aggressive signal types (no/yes), victim age class (adult vs. immature), and two interaction terms between victim age class and use of paedomorphic or aggressive signal types; see Table S3. Random intercepts were modelled to account for individual variation and repeated measures of victim and aggressor IDs. Affiliative signal types were excluded from all models as they were by default always used 100% after each aggression.

### Prediction C: Model 1.4

To test whether victims stopped signalling after having been consoled or reconciled, we fitted model 1.4 with the dependent variable of signalling persistence after receiving consolation / reconciliation (binary outcome of yes/no; Bernoulli distribution). The predictors in this model were aggression severity (mild/severe), piloerection during the bout (no/yes), reception of consolation (no/yes), victim age class (adult vs. immature), and interaction terms between victim age class and reception of consolation; see Table S3. We were unable to retain the variable *reconciliation* in our model due to limited sampling of data points for adults (i.e., in adults, only a total of five signal bouts were immediately followed by reconciliation, and of these, only one victim persisted once: LM, see Fig S3).

Random intercepts were modelled to account for individual variation and repeated measures of victim, aggressor IDs, and post-conflict ID (as some bouts were nested within the same conflict). Once again, affiliative signal types were excluded from all models as they were by default always used 100% after each aggression. Other than the other models which are being based on the level of the entire victim signalling display (in *N*=144 conflicts), this analysis is based on the level of signalling bouts (*N*=329).

### Predictions D-E: Models 1.5 and model 1.6

To test whether signal number changed with audience size and composition [number of audience members within 10m (D) and friends within 10 m (E)], we fitted a generalized mixed model with the dependent variable of signal number, represented as counts data (applying a negative binomial distribution, see Fig S2 model 1.5). The predictors in this model were aggression severity (mild/severe), piloerection (no/yes), number of audience members within 10 m (numeric), number of friends within 10m (counts), kin (i.e., mother) within 10 m (no/yes), and interaction terms between victim age class and audience members /friends /kin within 10 m; see Table S3. Random intercepts were modelled to account for individual variation and repeated measures of victim and aggressor IDs. Similarly, to test whether signal display duration (s) increases with audience members within 10 m (D) and friends within 10 m (E), we fitted a linear mixed model with a continuous dependent variable of signal display duration (applying a lognormal distribution, Fig S2 model 1.6). These two models include a smaller data set (N=142 conflicts), because data on audience composition was not present for three conflicts.

### Controlling for victim rearing

Our data were collected in a sanctuary environment where some individuals were orphans rescued from the bushmeat trade (see methods). Previous research on this population has revealed effects of rearing on bystander consolation, where mother-reared individuals are more likely to console victims than orphans; and orphans were slower to behaviourally recover than mother-reared, such as continuing to display distress signals after the conflict had ceased [47]. Therefore, we additionally controlled for the potential influence of victim rearing in all models.

However, these models had severe convergence issues, bad pareto k values and *Leave-one-out cross-validation* (LOOIC) also revealed that the fixed effect of victim rearing worsened model accuracies (Table S5). We thus excluded victim rearing from our presented models. Victim sex did not affect victim behaviour in previous a previous study [48] and was therefore not included in our models.

## Results

### Descriptive statistics

We analysed communicative signals produced by victims after *N*=144 aggressive attacks (mean = 5.3 attacks per victim, *SD* = 4.2) by conspecifics (“aggressors”). Fifty-three percent of such cases involved consolation as initiated by bystanders, 18% of cases involved reconciliation as initiated by aggressors, and 14% of cases involved renewed aggression by previous aggressors towards victims. Across conflicts (*N*=144), victims received severe aggression in 71%, or mild aggression (29%) of cases. The victims displayed piloerection (hair bristling from the body) in 36% of conflicts. The average victim signalling display duration across individuals was 63.8 s (*SD* = 79.5 s), which was 66.4 s (SD = 80.4 s) for adults and 62.8 s (SD = 79.6 s) for immatures.

### Component types

In N=144 conflicts, victims produced a total of N = 164 body signals (mean = 1.1, SD= 1.5 per individual/conflict), N = 165 gestures (mean = 1.1, SD =2.6), N = 548 facial expressions (mean = 3.8, SD = 3.7, additional *N* out-of-sight events = 13), and N = 901 vocalisations (mean = 6.3, SD = 6.2), see Fig. 1 A and Table S2.

**Fig. 1.**
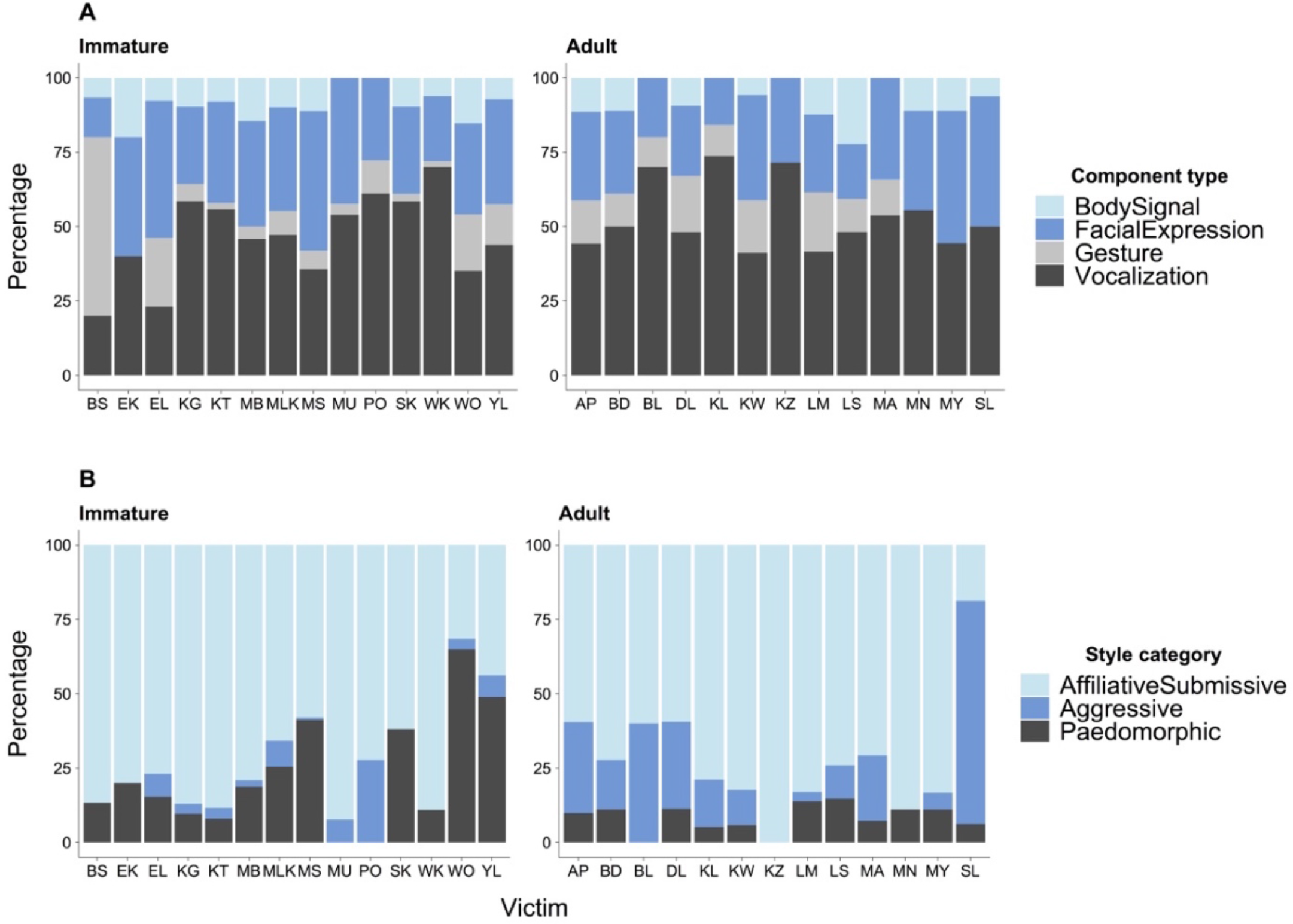
Percentage of signal components (A) and of communication style categories (B) used by victims in N= 144 aggressive conflicts, grouped by binary age class and victim ID.

### Communication styles

Across conflicts (N=144), victims most frequently produced *affiliative-submissive* signals (all individuals: mean = 72%, SD = 19%; immatures: mean = 72 %, SD = 19 %; adults: mean = 72 %, SD = 20 %), followed by *paedomorphic signals* (all individuals: mean = 16 %, SD = 16%; immatures: mean = 22 %, SD = 19 %; adults: mean = 9 %, SD = 5 %), and *aggressive signals* (all individuals: mean = 12 %, SD = 17 %; immatures: mean = 5 %, SD = 7 %; adults: mean = 20 %, SD = 21 %), see Fig. 1 B and Table S3.

#### A) Do paedomorphic signals increase the likelihood of receiving consolation and reconciliation, and reduce the risk of renewed aggression?

Controlling for aggression severity and victim arousal, victims were substantially more likely to be consoled by audience members after they produced paedomorphic signals following an aggressive attack (Fig. 2A and B; b = 1.57, SD = 0.77, 95% CredibIe Interval (“CrI”) [0.09, 3.12]; Table S4 model 1.1). There was only weak evidence however that paedomorphic signals increased the likelihood of reconciliation with aggressors (Fig. 2C and D; b = 2.55, SD = 1.75, 95% CrI [-0.52, 6.4]; Table S4 model 1.2), although this may be because the model’s predictive power was restricted due to small sample size of reconciliatory events in adults (Fig. S3 and Fig S2 model 1.2). There was also no evidence for any interaction effects between age and the use of paedomorphic signals with regards to the likelihood of being consoled or reconciled (Fig. 2A-D; Table S4 models 1.1 and 1.2). By contrast, adult victims were much less likely than immatures to receive renewed aggression when producing paedomorphic signals (Fig. 2E and F; b = 4.53, SD = 2.15, 95% CrI [0.73, 9.22]; Table S4 model 1.3).

**Fig. 2.**
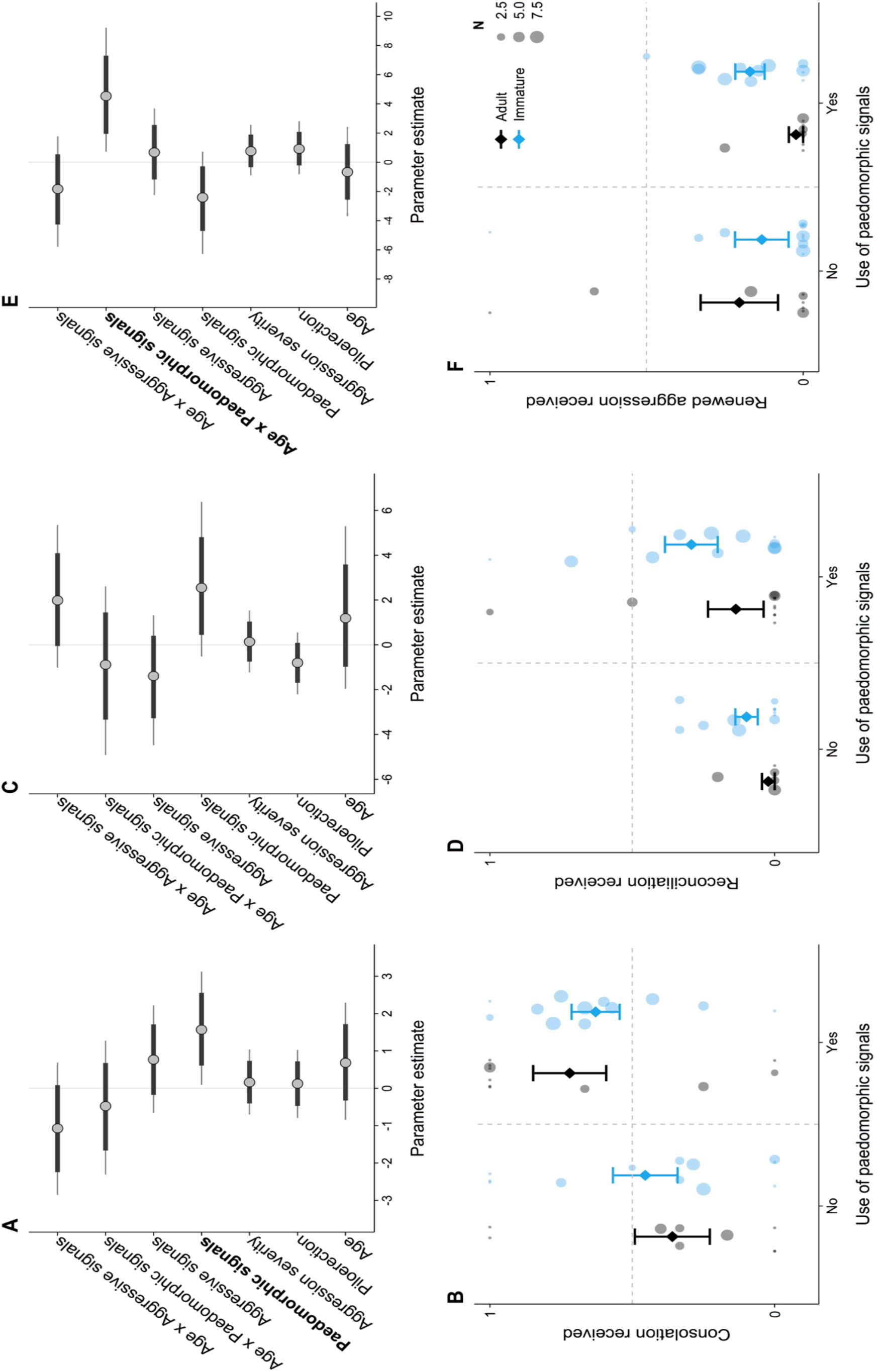
Uncertainty intervals from MCMC draws with all chains merged for models 1.1 (A), 1.2 (C) and 1.3 (E). Points denote posterior means, inner dark grey bands correspond to the 50% CrI, and the outer fine-lined bright grey bands correspond to the 95% CrI. Below are plots showing a summary of the raw data on the relationship between the proportion of having received consolation (B), reconciliation (D) or renewed aggression (F) in relation to paedomorphic signal use. Points denote proportions of victim consolation, reconciliation or renewed aggression received out of all observations of the victim, depending on whether or not these victims produced paedomorphic signals. Size of the points indicates the number of observations per victim. Diamonds depict mean proportion and upper and lower whiskers denote standard error of the mean.

Neither aggression severity nor piloerection had any clear impacts on the likelihood by which victims received consolation or renewed aggression, or reconciled with former opponents (Table S4 models 1.1-1.3).

#### B) Do aggressive signals reduce the likelihood of renewed aggression?

Controlling for aggression severity and victim arousal, the production of aggressive signals did not reduce the likelihood of renewed aggression (Fig. 2E; b = 0.68, SD = 1.5, 95% CrI [-2.25, 3.68]; Table S4 model 1.3), and there were no clear differences of this effect across age (Fig. 2E; b = -1.83, SD = 1.91, 95% CrI [-5.8, 1.78]; Table S4 model 1.3

#### C) Do victims stop signalling after having been consoled?

There was considerable variation in adult and immature victims’ signalling persistence depending on bystander behaviours. Adults were estimated to be less likely to persist in signalling after having been consoled as compared to immatures (Fig. 3A and B; b = 1.58, SD = 0.71, 95% CrI [0.21, 3.0], Table S4 model 1.4). There was also a greater likelihood of victims to persist in signalling when they were piloerect during a signalling bout (b = 0.85, SD = 0.43, 95% CrI [0.05, 1.74], Table S4 model 1.4). Aggression severity had no clear effect on signalling persistence (see Table S4 model 1.4).

**Fig. 3.**
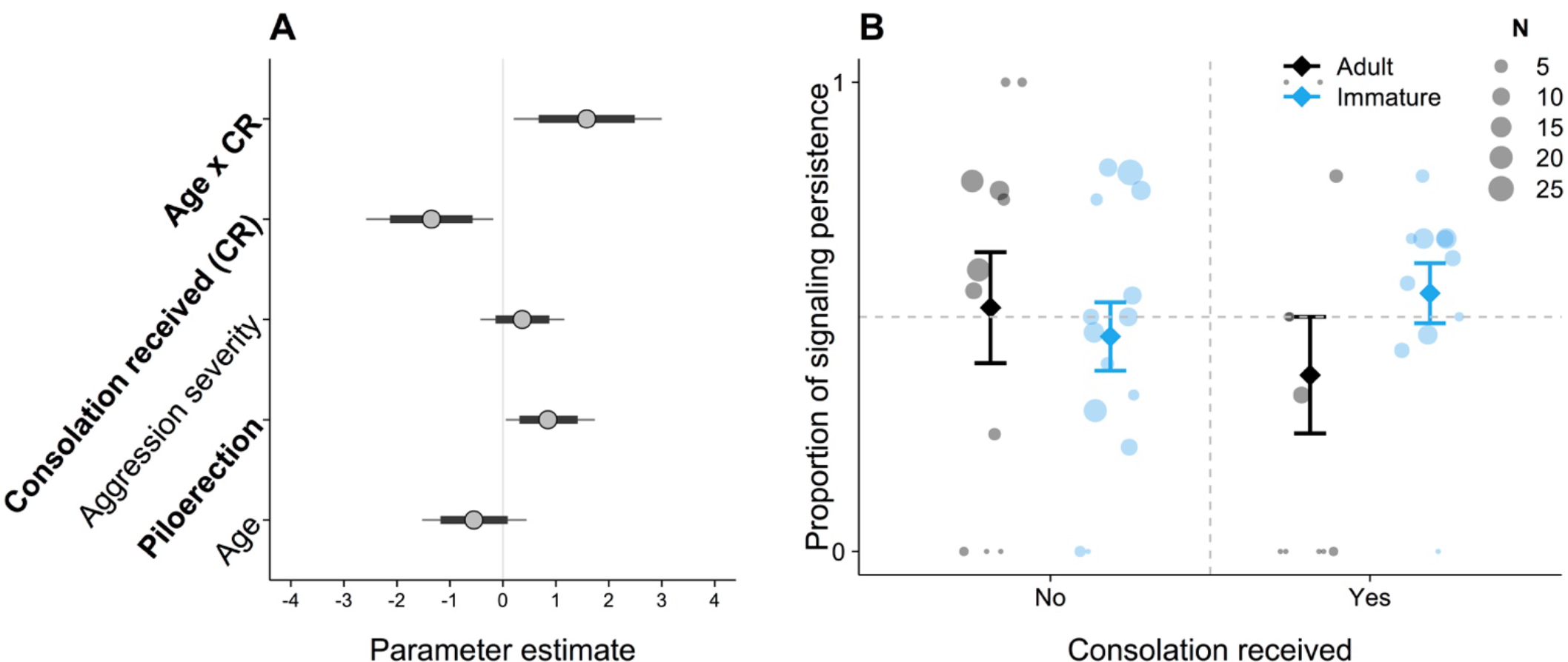
Uncertainty intervals from MCMC draws with all chains merged for model 1.4 (A). Points denote posterior means, inner dark grey bands correspond to the 50% CrI, and the outer fine-lined bright grey bands correspond to the 95% CrI. (B) shows a summary of the raw data on the relationship between the proportion of signalling persistence in relation to consolation. The size of the points indicates the number of observed bouts per victim. Diamonds depict mean proportion and upper and lower whiskers denote standard error of the mean.

#### D) Do victims increase signalling efforts when in presence of a larger audience?

When audience size increased, victims tended to produce slightly more signals (Fig. 4A and B; b = 0.04, SD = 0.04, 95% CrI [-0.03, 0.12], Table S4 model 1.5), with a weak interaction term between age and audience size, suggesting that immatures tend to produce more signals when in presence of a larger audience size compared to adults (Fig. 4A and B, b = 0.07, SD = 0.05, 95% CrI [-0.03, 0.17], Table S4 model 1.5).

**Fig. 4.**
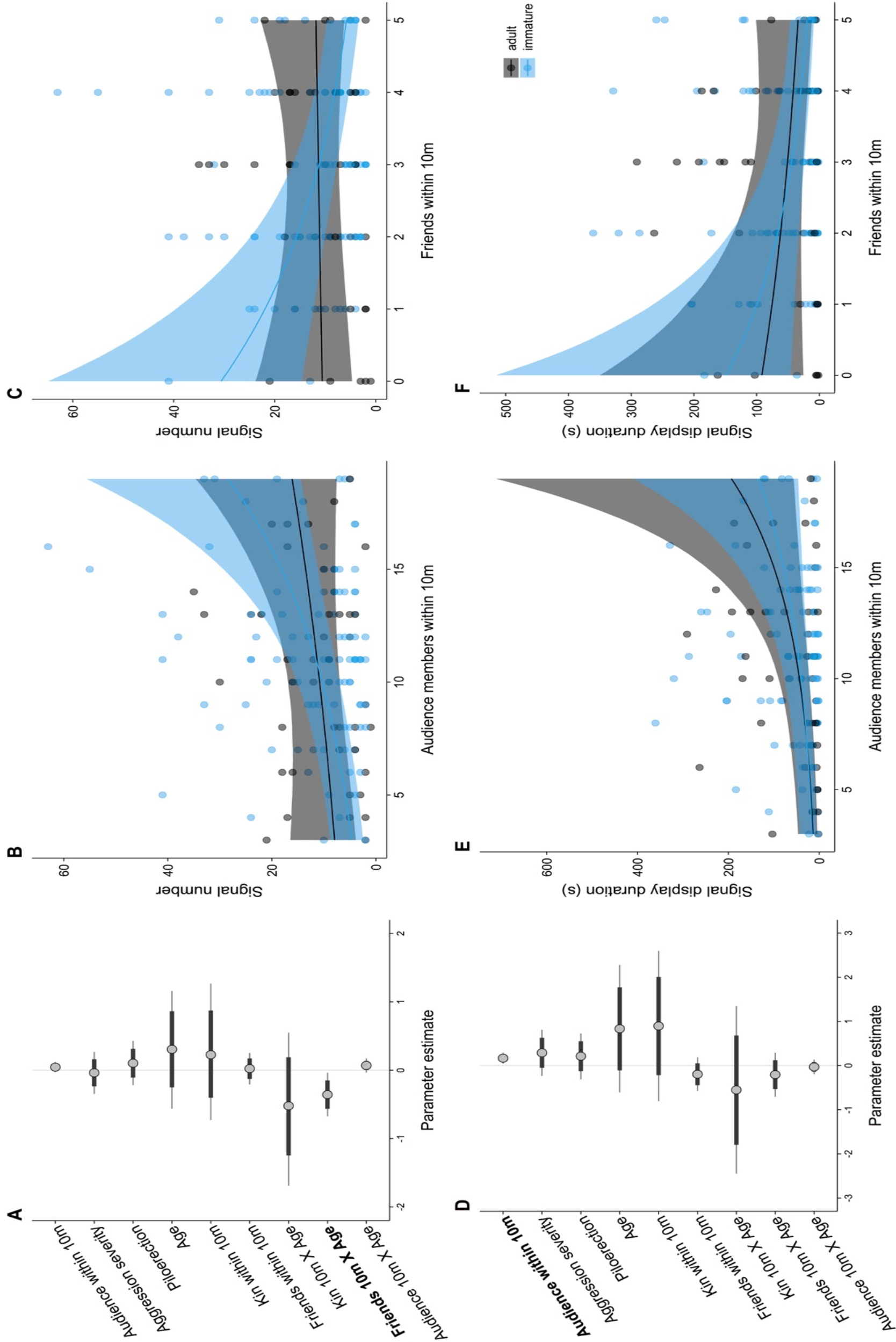
Uncertainty intervals from MCMC draws with all chains merged for model 1.5 (A-C) and 1.6 (D-F). Points denote posterior means, inner dark grey bands correspond to the 50% CrI, and the outer fine-lined bright grey bands correspond to the 95% CrI. Plots B, C, E and F represent a summary of the raw data on the relationships between signal number (B and C) and signal display duration (E and F) and audience members /friends within 10 m. Points denote single conflicts (i.e., one point represents signal number used and presence of an audience in one distinct conflict). Shaded upper and lower ribbon edges depict 95% credible intervals, and the mid-ribbon-line represent estimated posterior means

The main effect of audience became clearer when considering signal display duration: when audience size increased, victims (regardless of age) were more likely to engage in longer signal displays (Fig. 4D and E; b = 0.17, SD = 0.07, 95% CrI [0.04, 0.29], Table S4 model 1.6). These increased signalling efforts were not explained by arousal alone, because neither arousal cues (such as piloerection) nor severity of the aggression had any clear effect on signalling efforts (Fig. 4A and D, Table S4 models 1.5-1.6).

#### E) Do victims reduce signalling efforts when in presence of close social partners (friends)?

The model suggests that the number of friends within 10 m affected immature victim signal production, but not that of adults. Immatures produced less signals than adults when friends were nearby (Fig. 4A and C; b = -0.36, SD = 0.16, 95% CrI [-0.68, -0.04]; Table S4 model 1.5). This effect was less clear however for signal display duration, where no clear age differences nor general impact of friends could be determined (Fig. 4D and F; b = -0.21, SD = 0.25, 95% CrI [-0.71, 0.29]; Table S4 model 1.

## Discussion

We found that bonobo victims express emotional distress after aggressive attacks using a range of signalling techniques across modalities and that flexible adjustment of these emotional expressions influences receiver responses. Victims tailored their communicative signals depending on the audience size, as well as on bystanders’ identity and consolatory behaviour. Victim age also influenced signalling, whereby older victims were more likely to succeed in receiving a consolatory response when using paedomorphic signals compared to immature victims. Moreover, adults were more successful than immatures in preventing future aggression when using paedomorphic signals. Moreover, adults were appeared to be better able to suppress their distress signalling following consolation by bystanders, suggesting that a retainment of juvenile- like features has strategic functions in adulthood within the context of post-conflict interactions. However, as opposed to our prediction, the presence of close social partners of the victim in the audience had a clearer effect on signalling behaviour in immatures compared to adults, whereas kin generally had no clear influence on signalling behaviour, regardless of age.

There was notable variation in the ways victims expressed their distress, as shown by the range of signal components used (vocalisations, facial expressions, gestures, body signals) and the communication style deployed (aggressive, paedomorphic, affiliative- submissive). Traditionally, animal vocalisations and facial expressions have been perceived as emotional and involuntary [1], but recent research challenges this claim [20]: vocalisations can meet intentionality criteria if viewed in the framework used by gesture research [72] and vocalisations can actually be paired with other signal components that are by definition intentional: gestures and body signals [18, 19]. Our study supports the emerging view that great ape communicative displays during high-arousal contexts may concurrently be “intentional” and “emotional” [20]. This questions the purported dichotomy between emotional vocalisations or facial expressions on the one hand and intentional gestures or body signals on the other [1]. Such research challenges existing assumptions about the basis of cognitive control over emotional expressions in our primate relatives, and calls for a more inclusive analysis of emotion expressions in which signalling repertoire is looked at without assuming binary categorisation.

While our study suggests that victims can strategically deploy certain signals to promote prosocial responses in receivers, what remains nonetheless unclear is the extent to which these signals reliably *represent* the victims’ emotional states per se. Rather than being emotion signals of distress (like vocalisations and facial expressions presumably are), gestures and body signals might instead serve to specify certain intentions linked to the distressed state [1]. In other words, one might ask which type of emotion expressions reliably represents internal states? Are gestures and body signals honest indicators of victim emotional states, or are they communicative tools to win control over emotional episodes? Given that 25.7 % of conflicts in our sample were followed by vocalizations and/or facial expressions without any gestures or body signals, and given that gestures and body signals virtually never occurred without vocalization or facial expression, the latter assumption seems plausible. In humans this is different, since spoken language allows for the controlled expressions of emotion, endowing us to refer to and even explain our feelings, or to deceive and to influence receivers in tactic ways such as by reporting inaccurate states [73–75]; this is in line with prominent theories on language evolution [76, 77] suggesting that there was a shift from expressing intentions via gestures in nonhuman great apes and early hominins to expressing intent via mainly the vocal channel in modern humans. Future studies might evaluate this question on the basis of expressing affect in great apes, particularly in experimental settings, for instance by investigating whether apes would recognize others’ emotions by gestures or body signals alone [while removing facial or vocal cues, see 78 in preparation].

But how necessary is emotion signalling control, and to what extent is strategic emotional signalling human unique? Emotional control in humans enables us to engage in complex social life [79], and our capacity to adapt our emotion expressions to external circumstances may increase fitness. For instance, we might inhibit signals pertaining to our negative emotions, like anxiety in a job-interview, and instead voluntarily express signals of assertion or even dominance, in order to increase our chances of being selected [80]; we may otherwise exaggerate our disappointment or sadness to induce guilt in our receiver [81]. Likewise, losing emotional control can be detrimental; research shows that indeed, individuals who exhibit difficulties in emotion regulation have difficulties in navigating a healthy social life [82].

Although in human development, emotional regulation develops gradually and changes across the lifespan, infants already seem to have some control over their emotion states and expressions and may do so to alter the behaviour of recipients.

Louder and exaggerated crying in infancy, for instance, increases the chance of receiving caregivers’ attention and comfort, and this represents a crucial adaptation to increase infant survival [13, 14].

In our study, we found evidence of an apparent link between the production of expressions of distress and corresponding consolation by receivers, an expression of empathic concern. More specifically, we found that the likelihood of bystanders offering consolation (but not so for reconciliation) increased dramatically when bonobo victims (especially adults) used paedomorphic signals, i.e. those signals that rendered them more juvenile. In adults, the production of paedomorphic signals also reduced risk of renewed aggression by former opponents. Adults, compared to immatures, were less likely to receive renewed aggression by former opponents after having produced paedomorphic signals compared to when not. This suggests that, similar as in humans, emotion communication in bonobos may become more cognizant over the lifespan, with adults producing such signals more strategically. Additionally, it shows that paedomorphic signals may function in this species to elicit caring or prosocial responses in receivers, such as caregiving or consolation following aggression, which then, in turn, may have a calming effect on the victim (evidenced by the fact adult victims were more likely to stop signalling when having been consoled).

Evidence of paedomorphic victim signalling and its influence on bystander empathic tendencies in bonobos support broader evidence of the enhanced emotionality and juvenile nature of this species. Compared to other great apes, including chimpanzees, bonobos appear to have retained a range of paedomorphic features into adulthood (both morphologically and behaviourally), which some have suggested represent a positive selection towards friendliness and reduced aggression, akin to a process of self-domestication [46]. These findings thus makes bonobos a particularly interesting model to compare the evolutionary basis of emotional skills and communication in humans, as our own species has been thought to have undergone a process of self-domestication as well [83]. To address whether increased control over emotion expressions and the influence of paedomorphism was phylogenetically shared in our closest ancestor or whether it is a result of convergent self-domestication processes in bonobos and humans [83] cannot be decided here and should be tested by investigating such patterns in our other closest relatives, the chimpanzees.

More generally, evidence of enhanced consolatory responding by bystanders following paedomorphic victim signal highlights the close interplay between strategic distress signalling and corresponding empathic responding in receivers, which challenges the assumption that empathic responding is spontaneously offered [49]. Rather, it supports the view that empathy is elicited by certain communicative cues by distressed victims which may be under conscious control. An exciting future avenue would be to conduct further comparisons between humans and great apes to dissect potential other contexts beyond negative contexts in which controlled emotion communication may play a functional role, such as facilitating play or social affiliation [e.g., 73].

Evidence that adult victims were more likely than immatures to cease signalling after having been consoled supports the view that in bonobos, emotional signalling passes through a distinct developmental trajectory. As predicted, it suggests that adults express their emotions in more tactical ways – possibly to promote prosocial responding of the audience. This signalling technique may endow bonobos to confirm their status and ensure that consequent risks are low, or to alleviate their own stress level through peer support and affiliative contact. This level of awareness portrayed through the cessation of signal use following peer support did not reach the same levels in immatures. Instead, their signalling often continues even after consolatory events, suggesting that emotion signalling in immatures might involve less cognitive control and social awareness. Such developmental trajectories have also been detected in humans, where children only later, around the time when developing perspective taking skills, exhibit a more profound understanding of others’ emotion expressions and a capacity for more complex forms of empathy [85–87]. Our research shows that developmental pathways to emotion control also exist beyond our own species, which brings about exciting hypotheses for future research, related to how maternal style and early-life experiences might further shape emotionality in other animals [47, 48].

It is important to bear in mind though that piloerection also had a positive effect on signalling persistence, insofar as victims continued more often when being piloerect. This suggests that – in addition to bystanders’ consolatory contacts – the victims’ internal arousal level also played a role in signalling persistence and, crucially, that victims were honestly expressing distress states. Careful future analysis through experimental research using psycho-technological techniques like infrared thermography [88] should be used to understand the true extent to which arousal level drives signalling persistence in bonobo victims.

Likewise, one might argue that increased signal use and duration with greater audience size could be explained as a consequence of increased arousal levels, rather than cognitive control. Yet, our data in both models show that aggression severity and piloerection had no clear effect on signalling behaviour. Moreover, rather than being indiscriminate, we found that signalling was adapted to audience *composition* – especially in immatures: although immatures (like adults) communicated more when audience size increased, they produced substantially fewer signals when friends were close by (unlike adults). We predicted that adults should be more affected by audience composition, yet the opposite was true. What could this mean? One explanation could be that immatures are more vulnerable to aggression than adults, and hence are more in need of protection by friends (note that friends did not include kin, and that there was no effect of kin on signalling behaviour). This also means that bonobos seem to be aware of bystander relationships from an early age. The cognitive abilities involved cannot be ascertained with this data, and careful examination through experimental research is needed to further investigate this issue.

In sum, our current data – along with other findings regarding great ape emotion expressivity [see for a review 69] - alludes to the assumption that control over emotional expressions has a deep evolutionary history that in not human unique [2]. Across evolutionary time, the ecological expansion and progressively sophisticated social organization of early hominins suggest that they may have experienced a greater need to control their emotional outbursts both in ecological contexts, such as to reduce the risk of predation and increase hunting success, but also to facilitate more complex social interactions at dyadic, group and societal levels [2, 89]. Modern language then in turn might have further fuelled the expression of and ability to modify the expressions of one’s inner emotional state, favouring the emergence of complex cultural life today [74,75,90]. It is thus likely that some of the hallmarks of emotion control have preceded language evolution and have been present in our last common ancestor with the great apes [1].

We conclude that the current evidence from this study supports the theory that emotion control represents a highway to language evolution, with the building blocks of emotion control (i.e., tactic deployment of certain signals in the context of distress) likely already present in our last common ancestor with nonhuman great apes. This also complements other recent findings that other great ape species, notably chimpanzees [91, 92] and orangutans [29] can adjust their emotion expressions to their audience. Until now, previous research has mainly focused on single modalities, such as vocalisations [91, 92] and facial expressions [29] in isolation, lacking a multimodal view that considers how gestures and body signals may be used to add intentional components to such displays, and crucially, how such displays interact with receiver responses. Although we have made a start in this endeavour, further cross-modal and cross-component integration are required in order to assess the level of control involved in the emotion communication of our closest relatives. We advocate for more comparative data to improve insights into emotion control in other primate species as well as more distantly related species, by including all possible signal components. Such comparisons are crucial for shedding further light on the evolution of emotion capacities and their role in hominid evolution. Finally, our study informs on the mechanisms of empathy in our closest relatives, showing that empathy is not – as often presumed [see for a review 49] – a one-way process but rather an interplay between the victims’ (potentially strategic) communicative expressions and the bystanders response to those signals.

## Data availability statement

Video and audio files demonstrating communication in bonobo conflicts, as well as the data and R code supporting the article can be found in an online repository (figshare.com) under the private link:

## Supporting information

Electronic Supplementary Material

## Acknowledgements

We thank Frans de Waal for his support in the original studies from which these subsequent data are derived. We thank Pitshou Nsele Kayanga for assistance in data collection; Claudine André, Fanny Mehl, Fanny Minesi, Raphael Belais Dominique Morel and Pierrot Mbonzo and the Ministries of Research and Environment in the Democratic Republic of the Congo for their support (permit no. MIN.RS/SG/004/2009); the Lola ya Bonobo staff, particularly Stany Mokando and Jean-Claude Nzumbi. We thank Sally Street for statistical advice.

## Ethical statement

Ethical permission was received from ‘Les Amis des Bonobos du Congo’ (ABC) Scientific Committee and the Ministries of Research and Environment in the Democratic Republic of the Congo (permit no. MIN.RS/SG/004/2009), in compliance with all legal requirements for conducting research in DR Congo. This study also fully complied with Emory’s IACUC and Durham University’s guidelines for observational research with animals.

