## Supplementary material for "Tactical signalling by victims increases bystander consolation in bonobos": Electronic Supplementary Material

**Supplementary text 1**

To verify whether our coding was reliable, we recruited a second rater (ZU) to code a subset of the dataset, focusing on the number of signal types. ZU was untrained in ape behaviour and communication and coded signal types in 11% of the dataset (16 of 144 conflicts) following the ethogram in Table S2. She was blind to the coding sheet and asked to identify signal types independently (of all component possibilities such as gestures, vocalisations, facial expressions and body signals). The number of signal types was then compared; if signal types were coded differently (raters disagreed) this disagreement was recognized in the numbers. For instance, if RH coded *arm raise* twice but ZU did not code *arm raise* at all, then this would be noted as disagreement on gesture type *arm raise*.

**Table S1.** Summary of identity, number of observations, sex, age (continuous and binary), and rearing of victims (A) and aggressors (B).

**A. Victims**

| ID | N | Mean age 2011-2012, y | Binary age class | Sex | Rearing |
|---|---|---|---|---|---|
| AP | 11 | 12.0 | Adult | M | Orphaned |
| BD | 1 | 14.0 | Adult | F | Orphaned |
| BL | 3 | 10.3 | Adult | M | Orphaned |
| BS | 2 | 6.5 | Immature | M | Mother-reared |
| DL | 9 | 10.4 | Adult | M | Orphaned |
| EK | 1 | 6.0 | Immature | F | Mother-reared |
| EL | 2 | 9.0 | Immature | M | Orphaned |
| KG | 15 | 9.2 | Immature | M | Orphaned |
| KL | 2 | 13.0 | Adult | F | Orphaned |
| KT | 14 | 7.3 | Immature | F | Orphaned |
| KW | 1 | 13.0 | Adult | M | Orphaned |
| KZ | 1 | 22.0 | Adult | M | Orphaned |
| LM | 3 | 13.0 | Adult | M | Orphaned |
| LS | 4 | 10.5 | Adult | F | Mother-reared |
| MA | 4 | 11.0 | Adult | M | Orphaned |
| MB | 8 | 8.4 | Immature | M | Orphaned |
| MLK | 10 | 4.6 | Immature | F | Mother-reared |
| MN | 2 | 18.0 | Adult | M | Orphaned |
| MS | 9 | 5.3 | Immature | F | Orphaned |

| | | | | | |
|---|---|---|---|---|---|
| MU | 4 | 9.0 | Immature | F | Orphaned |
| MY | 1 | 19.0 | Adult | F | Orphaned |
| PO | 3 | 6.8 | Immature | M | Mother-reared |
| SK | 10 | 6.5 | Immature | F | Orphaned |
| SL | 1 | 14.0 | Adult | F | Orphaned |
| WK | 9 | 5.3 | Immature | F | Orphaned |
| WO | 8 | 3.8 | Immature | M | Mother-reared |
| YL | 7 | 7.9 | Immature | M | Orphaned |
| **Total** | **145** | **8.4** | - | - | - |

**B. Aggressors**

| ID | N | Mean age 2011-2012, y | Binary age class | Sex | Rearing |
|---|---|---|---|---|---|
| AP | 3 | 12.0 | Adult | M | Orphaned |
| BD | 11 | 14.0 | Adult | F | Orphaned |
| EK | 3 | 6.0 | Immature | F | Mother-reared |
| FZ | 22 | 13.0 | Adult | M | Orphaned |
| IB | 9 | 10.0 | Adult | M | Orphaned |
| KG | 3 | 9.2 | Immature | M | Orphaned |
| KL | 5 | 13.0 | Adult | F | Orphaned |
| KS | 12 | 13.0 | Adult | F | Orphaned |
| LI | 4 | 11.0 | Adult | F | Orphaned |
| LM | 4 | 13.0 | Adult | M | Orphaned |
| LS | 13 | 10.5 | Adult | F | Mother-reared |
| MA | 2 | 11.0 | Adult | M | Orphaned |
| MB | 2 | 8.4 | Immature | M | Orphaned |
| MD | 4 | 10.0 | Adult | M | Orphaned |
| MLK | 1 | 4.6 | Immature | F | Mother-reared |
| MN | 15 | 18.0 | Adult | M | Orphaned |
| MX | 1 | 26.0 | Adult | M | Orphaned |
| MY | 10 | 19.0 | Adult | F | Orphaned |
| OP | 8 | 17.0 | Adult | F | Orphaned |
| PO | 2 | 6.8 | Immature | M | Mother-reared |
| SL | 3 | 14.0 | Adult | F | Orphaned |

| | | | | | |
|---|---|---|---|---|---|
| SW | 7 | 15.0 | Adult | F | Orphaned |
| YL | 1 | 7.9 | Immature | M | Orphaned |
| **Total** | **145** | **13.4** | **-** | **-** | **-** |

**Table S2**. Frequency and definition of emotion signal components, signal types, and signal communication style categories (in parentheses, AS= affiliative submissive, P= paedomorphic, AG= aggressive) used by victims after aggressive attacks. Ethogram contains signal definitions as previously reported in the literature [16,30,50,92–98]. Signal components are listed according to the frequency of use (most common – rarest). All vocalisations can be heard as supplementary .wav files. Gesture or body signal types not demonstrated in photos here can either be seen under www.greatapedictionary.com, Genty et al. [93], Byrne et al. [30], or be requested by the lead author via.

| Facial expressions | N | Definition | Vocalisations | N | Definition | Gestures* | N | Definition | Body signals | N | Definition |
|---|---|---|---|---|---|---|---|---|---|---|---|
| Scream face(AS) 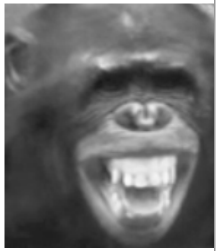 | 226 | The upper lips are raised and the lip corners pulled back, while exposing the upper teeth. Additionally, the lower lip is depressed and subject exposed the lower teeth, with the mouth being stretched wide open and the lips parted. | Peep shriek/scream/yelp (AS) | 426 | Sounds shriller than the other peeps and is of longer duration. A pure, tonal unmodulated sound with no or few harmonics. Facial expression always involves partial or complete showing of teeth. | Hand reach(P) | 34 | Holding a hand toward another individual by extending the arm and hand. | Tantrum position(P) 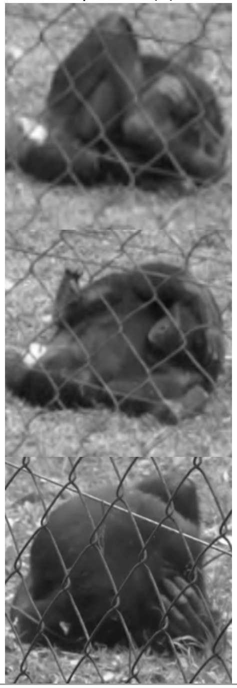 | 28 | Haunching close to the ground with the entire body, as if frozen, can be either with back on the ground and stomach exposed, or while lying on the stomach; signalers often look at the aggressor during the display, while the head is interchangeably protected by arms/hands (as if signaler was hiding). |

| | | | | | | | | | | | |
|---|---|---|---|---|---|---|---|---|---|---|---|
| Bared teeth(AS) 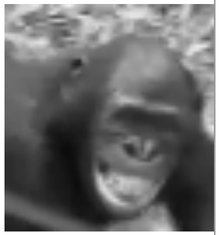 | 166 | Retraction of the lips resulting in partial or complete exposure of the teeth and gums, with mouth practically closed and often silent, without vocalisation. Displayed in agonistic and approach-retreat interactions, during sexual interactions. Performed by both subordinate and dominant individuals, expression of fear, nervousness, or hesitation. Demonstrates non-hostile intentions, function as appeasement and reassurance signal. | Victim scream (AS) | 234 | High-pitched, with shrill and rasping sounds given at full vocal strength, large number of harmonics and long in duration, accompanied by teeth-baring (complete lip retraction, exposing both teeth and gums, the mouth may be wide open). | Touch(AS) | 19 | Touching gently another individual's body part with palm of hand, for under 2s. | Concave back present(AS) | 26 | Sitting in front of recipient with arched back to expose genitals with legs spread apart, sexual invitation. |
| Pout face(P) 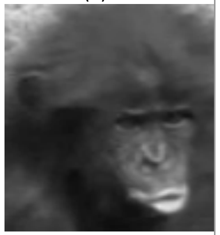 | 116 | Lips are pursed forward and curled outward in front resulting in a circular opening. The lips are pressed together at the mouthcorners. Displayed to request for food or request physical contact. | Pout moan (P) | 79 | Low-pitched, melodious call sounding like a whining "hoo-hoo", accompanied by pout face (lips are pursed forward and curled outward in front resulting in circular opening). Used primarily by infants for requests, used by adults during sexual | Arm raise(P) | 14 | Raising one arm above the head. | Exaggerated concave back present(AS) | 19 | Standing quadrupedally in front of recipient with ventral side up to expose genitals with legs spread apart. |

| | | | | | | | | | | | |
|---|---|---|---|---|---|---|---|---|---|---|---|
| | | Sometimes displayed during grooming. Can express disappointment. | | | initiations, food begging, or as a appeasement signal in agonistic interactions. | | | | | | |
| Threat face(AG) 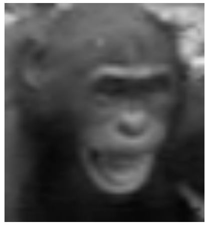 | 20 | Mouth opened with intense stare at opponent; head often tilted downwards, teeth can be visible. Often seen during production threat barks or contest hoot vocalisations. Displayed in aggressive chases towards others. | Pout whimper (P) | 48 | Also known as “whimper”. Longer voicing than pout moan, often long lasting and connected with severe emotional distress. | Throw object(AG) | 10 | Throwing an object in direction of another individual. | Tantrum bounce(P) | 14 | Bouncing up and down on a vertical axis with upper body (often combined with victim screams and bared-teeth) |
| Grin low closed(P) 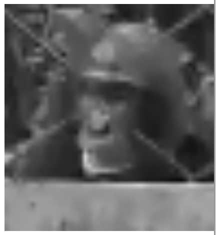 | 11 | Similar to bared-teeth but only the lower of the anterior teeth are revealed, often follows whimpering and precedes crying. Linked with whimpering, distress, fear. | Threat bark (AG) | 27 | A shrill bark used during aggressive attacks, or if bothered by another individual approach or behavior. | Stomp(AG) | 10 | Stamping the ground forcefully with sole of foot. | Pirouette(P) | 14 | Twirling movement of whole body around the body y axis |
| Tense mouth/ bulging lip face (AG) 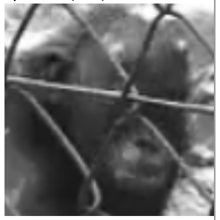 | 5 | Lips in horizontal tension and drawn inward. No vocalisations accompanying this expression. | Peep(AS) | 39 | High-frequency, closed mouth vocalisation , short in duration and characterised by a simple, flat acoustic form composed of several harmonics that are generally un-modulated. Produced across a large array of | Gentle grab(P) | 8 | Grabbing gently another individual’s body part with closed hand(s). | Bipedal present(AS) | 12 | Standing bipedally in front of recipient with arms spread apart, sexual invitation. |

| | | | | | | | | | | | |
|---|---|---|---|---|---|---|---|---|---|---|---|
| | | | | | contexts, especially common during feeding events and travel, but also during agonistic contexts. | | | | | | |
| Horizontal pout face(P) 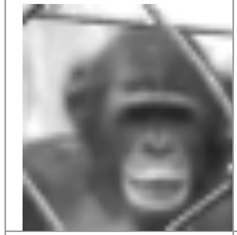 | 2 | Similar to grin low closed but upper lip slightly opens, so that anterior upper and lower teeth show. Always connected with pour whimpering and crying. Linked with distress, fear. | Agonistic peep(AG) | 25 | Similar to peep but louder and used in aggressive threat to others. | Rough grab(AG) | 8 | Grabbing aggressively and roughly another individual's body part with closed hand(s); often in fights. | Rump present(AS) | 10 | Standing quadrupedally in front of recipient with dorsal side up to expose hindquarters, while looking back at recipient, sexual invitation. |
| Relaxed playface(AS) 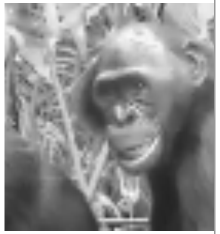 | 2 | Relaxed open mouth, only lower teeth are showing. Facial expression accompanies social play in non-aggressive contexts. | Contest hoot(AG) | 22 | Call sequences consisting of an introductory phase (modulated inverted u-shape form), an escalation phase composed of several stereotyped units (unmodulated inverted u-shape), and a let-down phase. Used by males to challenge another individual and display status to others. | Slap(AG) | 8 | Slapping forcefully and singly another individual with palm of hand. | Tantrum role(P) | 8 | Rolling back and forth on the ground. |
| Full playface(AS) 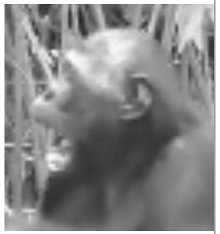 | 1 | Full open mouth, upper and lower teeth are showing. Facial expression accompanies social play in non-aggressive contexts. | Unknown | 2 | Unknown vocalisation | Hand(s) on(AS) | 7 | Touching head (or other body part) of another individual with palm(s) of hand(s) or feet and maintaining touch for more than 2 s. | Body shake(AS) | 4 | Signaller shakes entire body from shoulder joints, while looking at recipient from a distance. Often paired with bared-teeth display. |

| | | | | | | | | | | | |
|---|---|---|---|---|---|---|---|---|---|---|---|
| | | | | | | Head shake(AS) | 7 | Shaking head from side to side on horizontal axis. | Gallop(AG) | 4 | Exaggerated running with forelegs playfully stamping the floor (similar to a child imitating a horse galloping). |
| | | | | | | Stretch over(AS) | 7 | Stretching and raising arm till about head level with the palm facing downwards, sexual invitation. | Object dragging(AG) | 4 | Dragging object held in hand alongside of the body (usually branch) while moving forward, charging display. |
| | | | | | | Slap object(AG) | 6 | Slapping forcefully and singly object with palm of hand. | Bite(AG) | 3 | Recipient's body is held between or against lips or teeth of signaller |
| | | | | | | Embrace(P) | 5 | Signaller wraps arm(s) around recipient and maintains physical contact. | Look/peering(AS) | 3 | Signaller holds eye contact with recipient lasting >2 s |
| | | | | | | Hand shake(P) | 4 | Shaking hand vigorously and loosely from wrist joint. | Somersault(P) | 3 | Twirling forward movement of whole body around the body x axis. |
| | | | | | | Head nod(AS) | 3 | Nodding head up and down in the body x axis. | Ventral present(AS) | 3 | Lying on back in front of recipient with legs spread apart to expose genitals. |
| | | | | | | Leg reach(AS) | 3 | Holding a foot toward another individual by extending the leg and foot | Climb on(P) | 3 | Young individual touching mother's back from behind and making an attempt to climb on (often with one leg half on the mother's back) and staying in that position (but not actually climbing on her back). |

| | | | | | | | | | | | |
|---|---|---|---|---|---|---|---|---|---|---|---|
| | | | | | | Punch(AG) | 2 | Hitting another individual forcefully and singly with fist(s) or wrist(S). | Bipedal swagger(AG) | 1 | Lateral swaying of the upper body. |
| | | | | | | Push(AG) | 2 | Pushing away another individual with hand(s) or arm(s). | Chase threat(AG) | 1 | Signaller running towards another individual as though chasing the individual, often combined with facial threat expression. |
| | | | | | | Arm swing(AG) | 1 | Swinging arm(s) back and forth on side, either once or repetitively. | Dangle(P) | 1 | Signaller hangs from arm(s) above another, may shake feet/legs, typically audible with movement in canopy |
| | | | | | | Grab pull(P) | 1 | Grabbing gently another individual's body part with closed hand(s) and pulling towards self. | Head stand (P) | 1 | Signaller bends forward and places head on ground |
| | | | | | | Hand wave off(AS) | 1 | Raising arm(s) and waving it away from self. | Stiff walk(AG) | 1 | Walking with rigid forelegs, with a slow exaggerated movement. |
| | | | | | | Leg up(AS) | 1 | Raising one leg at hip level. | Tandem walk(P) | 1 | Subject positions arm over the body of the recipient and both walk forward while maintaining position |
| | | | | | | Rhythmic stomp(AG) | 1 | Stamping the ground alternatively with one foot then the other very rapidly. | | | |
| | | | | | | Shoo/ hand(s) fling(AG) | 1 | Rapid movement of | | | |

| | | | | | | | | |
|---|---|---|---|---|---|---|---|---|
| | | | | | | | | hand(s) or arm(s) from the signaller towards the recipient |
| | | | | | | Stroke(AS) | 1 | Stroking another individual with gentle back and forth movement of palm of hand or fingers. |
| | | | | | | Tap(AS) | 1 | Tapping repetitively another individual with palm of hand, with firm short contact of the fingers to the signaler's body (may include rhythmic repetition). |

**Table S3**. Communication styles categories (with total N of signals and percentage use of communication style categories) across victims. P= Paedomorphic signals; AS= Affiliative submissive signals; AG= Aggressive signals.

| ID | Number P | Number AS | Number AG | Age class | Total N signals used | % P | % AS | % AG |
|---|---|---|---|---|---|---|---|---|
| **AP** | 13 | 77 | 40 | Adult | 130 | 10.0 | 59.2 | 30.8 |
| **BD** | 2 | 13 | 3 | Adult | 18 | 11.1 | 72.2 | 16.7 |
| **BL** | 0 | 12 | 8 | Adult | 20 | 0.0 | 60.0 | 40.0 |
| **BS** | 2 | 13 | 0 | Immature | 15 | 13.3 | 86.7 | 0.0 |
| **DL** | 12 | 64 | 31 | Adult | 107 | 11.2 | 59.8 | 29.0 |
| **EK** | 1 | 4 | 0 | Immature | 5 | 20.0 | 80.0 | 0.0 |
| **EL** | 2 | 10 | 1 | Immature | 13 | 15.4 | 76.9 | 7.7 |
| **KG** | 15 | 134 | 5 | Immature | 154 | 9.7 | 87.0 | 3.2 |
| **KL** | 1 | 15 | 3 | Adult | 19 | 5.3 | 78.9 | 15.8 |
| **KT** | 8 | 117 | 5 | Immature | 130 | 6.2 | 90.0 | 3.8 |
| **KW** | 1 | 14 | 2 | Adult | 17 | 5.9 | 82.4 | 11.8 |
| **KZ** | 0 | 7 | 0 | Adult | 7 | 0.0 | 100.0 | 0.0 |
| **LM** | 9 | 54 | 2 | Adult | 65 | 13.8 | 83.1 | 3.1 |
| **LS** | 4 | 20 | 3 | Adult | 27 | 14.8 | 74.1 | 11.1 |
| **MA** | 4 | 29 | 8 | Adult | 41 | 9.8 | 70.7 | 19.5 |
| **MB** | 9 | 38 | 1 | Immature | 48 | 18.8 | 79.2 | 2.1 |
| **MLK** | 41 | 106 | 14 | Immature | 161 | 25.5 | 65.8 | 8.7 |
| **MN** | 1 | 8 | 0 | Adult | 9 | 11.1 | 88.9 | 0.0 |
| **MS** | 61 | 83 | 1 | Immature | 145 | 42.1 | 57.2 | 0.7 |
| **MU** | 0 | 24 | 2 | Immature | 26 | 0.0 | 92.3 | 7.7 |
| **MY** | 2 | 15 | 1 | Adult | 18 | 11.1 | 83.3 | 5.6 |
| **PO** | 0 | 13 | 5 | Immature | 18 | 0.0 | 72.2 | 27.8 |
| **SK** | 44 | 70 | 0 | Immature | 114 | 38.6 | 61.4 | 0.0 |
| **SL** | 1 | 3 | 12 | Adult | 16 | 6.3 | 18.8 | 75.0 |
| **WK** | 22 | 187 | 0 | Immature | 209 | 10.5 | 89.5 | 0.0 |
| **WO** | 72 | 35 | 4 | Immature | 111 | 64.9 | 31.5 | 3.6 |
| **YL** | 68 | 60 | 10 | Immature | 138 | 49.3 | 43.5 | 7.2 |

**Table S4.** Summary of Bayesian regression models. Reference levels are marked in parentheses [ ].

| **Model 1.1 Consolation (conflicts, N=144)** | | | | | | | |
|---|---|---|---|---|---|---|---|
| | eff. *N* | $\hat{R}$ | *b* | *S.D.* | 2.50% | 97.50% | Default Prior |
| Intercept | 26823 | 1 | -1.19 | 0.75 | -2.7 | 0.25 | Student *t* (3,0,10) |
| Aggression severity [mild] | 32000 | 1 | 0.16 | 0.44 | -0.7 | 1.04 | Student *t* (3,0,10) |
| Piloerection [no] | 32000 | 1 | 0.12 | 0.47 | -0.79 | 1.03 | Student *t* (3,0,10) |
| Paedomorphic signals [no] | 26476 | 1 | 1.57 | 0.77 | 0.09 | 3.12 | Student *t* (3,0,10) |
| Age [adult] | 24711 | 1 | 0.68 | 0.8 | -0.84 | 2.29 | Student *t* (3,0,10) |
| Aggressive signals [no] | 28095 | 1 | 0.77 | 0.74 | -0.66 | 2.22 | Student *t* (3,0,10) |
| Age [adult] x Paedomorphic signals [no] | 25981 | 1 | -0.48 | 0.91 | -2.31 | 1.28 | Student *t* (3,0,10) |
| Age [adult] x Aggressive signals [no] | 27547 | 1 | -1.07 | 0.91 | -2.86 | 0.69 | Student *t* (3,0,10) |
| Aggressor Intercept | 10776 | 1 | 0.48 | 0.34 | 0.02 | 1.29 | Student *t* (3,0,10) |
| Victim Intercept | 9520 | 1 | 0.53 | 0.37 | 0.02 | 1.42 | Student *t* (3,0,10) |
| **Model 1.2 Reconciliation (conflicts, N=144)** | | | | | | | |
| | eff. *N* | $\hat{R}$ | *b* | *S.D.* | 2.50% | 97.50% | Default Prior |
| Intercept | 11521 | 1 | -4.31 | 1.82 | -8.5 | -1.35 | Student *t* (3,0,10) |
| Aggression severity [mild] | 32000 | 1 | 0.13 | 0.7 | -1.23 | 1.53 | Student *t* (3,0,10) |
| Piloerection [no] | 32000 | 1 | -0.8 | 0.7 | -2.2 | 0.55 | Student *t* (3,0,10) |
| Paedomorphic signals [no] | 12506 | 1 | 2.55 | 1.75 | -0.52 | 6.38 | Student *t* (3,0,10) |
| Age [adult] | 12531 | 1 | 1.19 | 1.85 | -1.96 | 5.29 | Student *t* (3,0,10) |
| Aggressive signals [no] | 17757 | 1 | -1.39 | 1.46 | -4.48 | 1.32 | Student *t* (3,0,10) |
| Age [adult] x Paedomorphic signals [no] | 12958 | 1 | -0.89 | 1.91 | -4.92 | 2.61 | Student *t* (3,0,10) |
| Age [adult] x Aggressive signals [no] | 17611 | 1 | 1.99 | 1.63 | -1.02 | 5.35 | Student *t* (3,0,10) |
| Aggressor Intercept | 8473 | 1 | 1.5 | 0.74 | 0.3 | 3.23 | Student *t* (3,0,10) |
| Victim Intercept | 7829 | 1 | 1.21 | 0.73 | 0.09 | 2.92 | Student *t* (3,0,10) |
| **Model 1.3 Renewed aggression (conflicts, N=144)** | | | | | | | |
| | eff. *N* | $\hat{R}$ | *b* | *S.D.* | 2.50% | 97.50% | Default Prior |
| Intercept | 9239 | 1 | -4.75 | 2.13 | -9.79 | -1.42 | Student *t* (3,0,10) |
| Aggression severity [mild] | 32000 | 1 | 0.76 | 0.88 | -0.9 | 2.57 | Student *t* (3,0,10) |
| Piloerection [no] | 32000 | 1 | 0.92 | 0.91 | -0.82 | 2.81 | Student *t* (3,0,10) |
| Paedomorphic signals [no] | 18244 | 1 | -2.41 | 1.78 | -6.28 | 0.73 | Student *t* (3,0,10) |
| Age [adult] | 18240 | 1 | -0.67 | 1.55 | -3.7 | 2.42 | Student *t* (3,0,10) |
| Aggressive signals [no] | 20159 | 1 | 0.68 | 1.49 | -2.25 | 3.68 | Student *t* (3,0,10) |
| Age [adult] x Paedomorphic signals [no] | 14919 | 1 | 4.53 | 2.15 | 0.73 | 9.22 | Student *t* (3,0,10) |
| Age [adult] x Aggressive signals [no] | 18482 | 1 | -1.83 | 1.91 | -5.8 | 1.78 | Student *t* (3,0,10) |
| Aggressor Intercept | 7402 | 1 | 3.09 | 1.45 | 1.14 | 6.73 | Student *t* (3,0,10) |
| Victim Intercept | 6500 | 1 | 1.2 | 0.92 | 0.06 | 3.45 | Student *t* (3,0,10) |

| **Model 1.4 Signalling persistence (signalling bouts, N=329)** | | | | | | | |
|---|---|---|---|---|---|---|---|
| | n_eff | Rhat | mean | sd | 2.50% | 97.50% | Default Prior |
| Intercept | 16155 | 1 | -0.17 | 0.54 | -1.3 | 0.83 | Student *t* (3,0,10) |
| Aggression severity [mild] | 32000 | 1 | 0.36 | 0.4 | -0.42 | 1.16 | Student *t* (3,0,10) |
| Piloerection [no] | 13049 | 1 | 0.85 | 0.43 | 0.05 | 1.74 | Student *t* (3,0,10) |
| Consolation received [no] | 22619 | 1 | -1.35 | 0.61 | -2.58 | -0.18 | Student *t* (3,0,10) |
| Age [adult] | 23058 | 1 | -0.55 | 0.5 | -1.52 | 0.45 | Student *t* (3,0,10) |
| Age [adult] x Consolation received [no] | 22775 | 1 | 1.58 | 0.71 | 0.21 | 3 | Student *t* (3,0,10) |
| Aggressor Intercept | 9381 | 1 | 0.34 | 0.27 | 0.01 | 0.99 | Student *t* (3,0,10) |
| Victim Intercept | 4922 | 1 | 1.09 | 0.34 | 0.4 | 1.78 | Student *t* (3,0,10) |

| **Model 1.5 Signal number (conflicts, N=142)** | | | | | | | |
|---|---|---|---|---|---|---|---|
| | eff. *N* | $\hat{R}$ | *b* | *S.D.* | 2.50% | 97.50% | Default Prior |
| Intercept | 32000 | 1 | 1.87 | 0.37 | 1.15 | 2.61 | Student *t* (3,2,10) |
| Aggression severity [mild] | 32000 | 1 | -0.04 | 0.16 | -0.35 | 0.27 | Student *t* (3,2,10) |
| Piloerection [no] | 32000 | 1 | 0.1 | 0.16 | -0.22 | 0.43 | Student *t* (3,2,10) |
| Audience within 10 m | 17640 | 1 | 0.04 | 0.04 | -0.03 | 0.12 | Student *t* (3,2,10) |
| Age [adult] | 32000 | 1 | 0.3 | 0.44 | -0.56 | 1.16 | Student *t* (3,2,10) |
| Friends within 10 m | 17601 | 1 | 0.02 | 0.12 | -0.21 | 0.25 | Student *t* (3,2,10) |
| Kin within 10 m | 25688 | 1 | 0.23 | 0.51 | -0.73 | 1.27 | Student *t* (3,2,10) |
| Age [adult] x Audience within 10 m | 15429 | 1 | 0.07 | 0.05 | -0.03 | 0.17 | Student *t* (3,2,10) |
| Age [adult] x Friends within 10 m | 15289 | 1 | -0.36 | 0.16 | -0.68 | -0.04 | Student *t* (3,2,10) |
| Age [adult] x Kin within 10 m | 24031 | 1 | -0.52 | 0.57 | -1.69 | 0.55 | Student *t* (3,2,10) |
| Aggressor Intercept | 7641 | 1 | 0.27 | 0.13 | 0.03 | 0.54 | Student *t* (3,0,10) |
| Victim Intercept | 7978 | 1 | 0.32 | 0.13 | 0.07 | 0.59 | Student *t* (3,0,10) |

| **Model 1.6 Signal display duration (conflicts, N=142)** | | | | | | | |
|---|---|---|---|---|---|---|---|
| | eff. *N* | $\hat{R}$ | *b* | *S.D.* | 2.50% | 97.50% | Default Prior |
| Intercept | 32000 | 1 | 1.81 | 0.61 | 0.62 | 3 | Student *t* (3,3,10) |
| Aggression severity [mild] | 32000 | 1 | 0.29 | 0.27 | -0.23 | 0.81 | Student *t* (3,3,10) |
| Piloerection [no] | 32000 | 1 | 0.21 | 0.26 | -0.31 | 0.73 | Student *t* (3,3,10) |
| Audience within 10 m | 19512 | 1 | 0.17 | 0.07 | 0.04 | 0.29 | Student *t* (3,3,10) |
| Age [adult] | 32000 | 1 | 0.83 | 0.73 | -0.61 | 2.28 | Student *t* (3,3,10) |
| Friends within 10 m | 20373 | 1 | -0.2 | 0.19 | -0.58 | 0.18 | Student *t* (3,3,10) |
| Kin within 10 m | 24950 | 1 | 0.9 | 0.87 | -0.81 | 2.59 | Student *t* (3,3,10) |
| Age [adult] x Audience within 10 m | 18635 | 1 | -0.03 | 0.08 | -0.2 | 0.13 | Student *t* (3,3,10) |
| Age [adult] x Friends within 10 m | 19581 | 1 | -0.21 | 0.25 | -0.71 | 0.29 | Student *t* (3,3,10) |
| Age [adult] x Kin within 10 m | 23558 | 1 | -0.55 | 0.97 | -2.45 | 1.35 | Student *t* (3,3,10) |
| Aggressor Intercept | 13234 | 1 | 0.22 | 0.16 | 0.01 | 0.61 | Student *t* (3,0,10) |
| Victim Intercept | 9255 | 1 | 0.39 | 0.2 | 0.03 | 0.82 | Student *t* (3,0,10) |

* **Abbreviations**: *b*= Estimated mean of the posterior distribution; *S.D.*= Standard deviation of the posterior distribution; 2.50% and 97.5%= Two-sided 95% Credible intervals based on quantiles; $\hat{R}$=R hat value, provides information about the convergence of the MCMC algorithm - if larger than 1.1, chains have not converged and model is not accurate; eff. *N*= number of effective sample size (i.e., number of independent samples from the posterior distribution, which would be expected to give the same standard error of posterior mean as obtained from dependent samples returned by MCMC algorithms).

**Table S5**. Leave-one-out cross-validation (LOOIC) comparing model fit including or excluding the variable “victim rearing” [99]. The first listed model version (“reduced”: excluding victim rearing) is compared against secondary model versions (“full”: including victim rearing). The difference in the expected log predictive densities (ELPD) were negative, suggesting that reduced models are preferred (i.e., they had lower LOOIC values and better convergence). Victim rearing is thus not expected to improve model accuracy, for which reason only the reduced models excluding victim rearing are presented in the main paper. For problematic Pareto k values, we set IC comparisons to RELOO=TRUE.

| **Model** | **ELPD difference** | **SE difference** |
|---|---|---|
| Model 1.1 (reduced) | -1.1 | 0.9 |
| Model 1.1 (full) | | |
| Model 1.2 (reduced) | -1.3 | 1.1 |
| Model 1.2 (full) | | |
| Model 1.3 (reduced) | -3.9 | 1.4 |
| Model 1.3 (full) | | |
| Model 1.4 (reduced) | -1.4 | 0.5 |
| Model 1.4 (full) | | |
| Model 1.5 (reduced) | -23.0 | 2.8 |
| Model 1.5 (full) – divergent chains | | |
| Model 1.6 (reduced) | -28.5 | 1.2 |
| Model 1.6 (full) – divergent chains | | |

**Fig S1.** Trace plots of MCMC chains and posterior distributions of all models.

Model 1.1

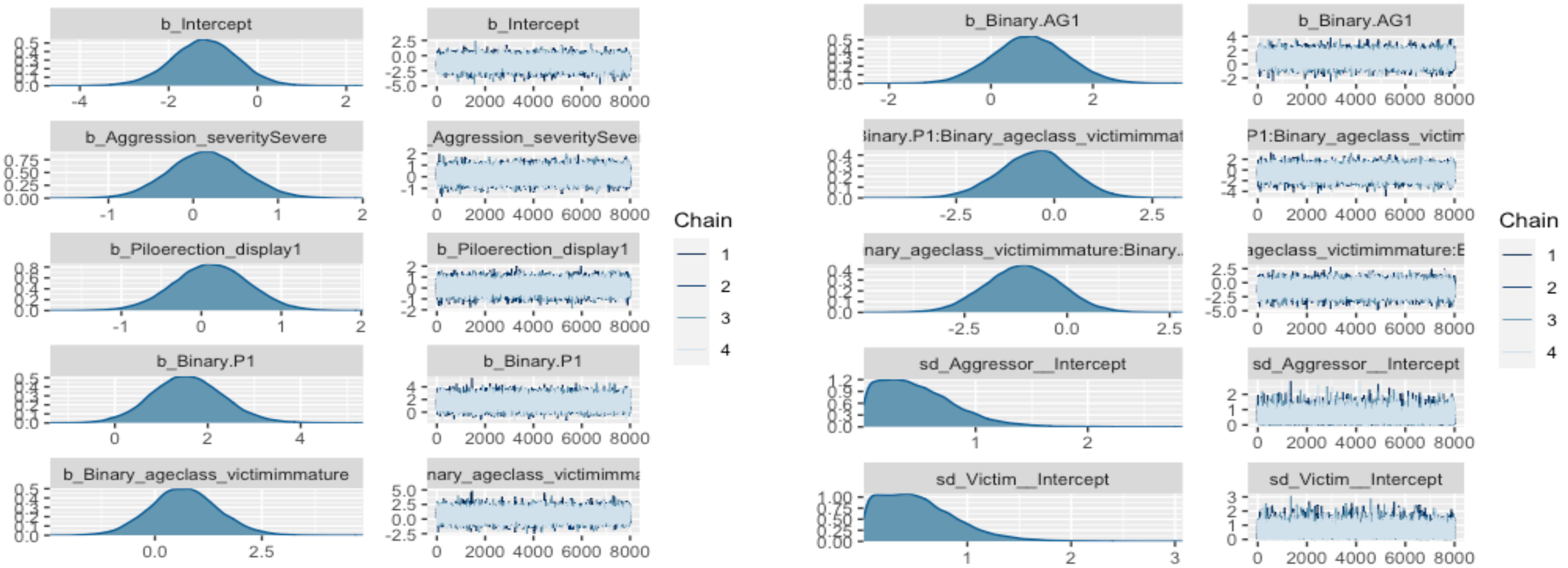

Model 1.2

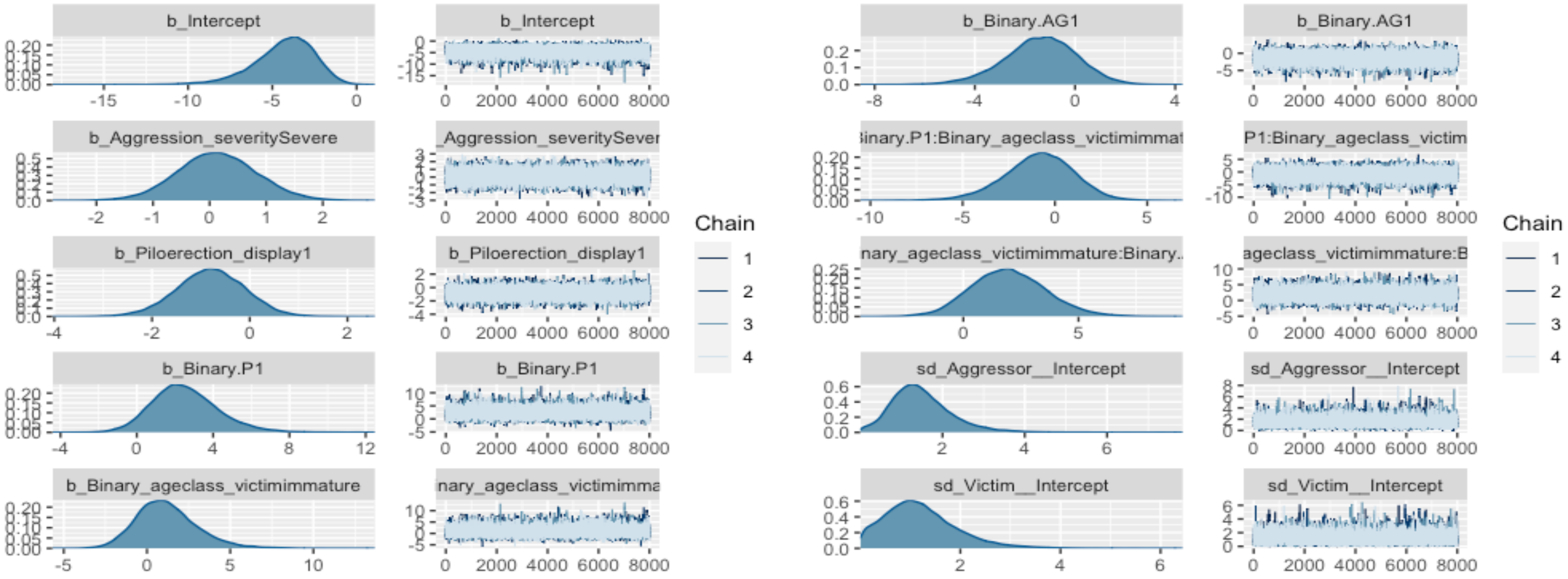
b_Intercept
b_Intercept
b_Binary.AG1
b_Binary.AG1
b_Aggression_severitySevere
Aggression_severitySever
inary.P1:Binary_ageclass_victimimmat
P1:Binary_ageclass_victim
b_Piloerection_display1
b_Piloerection_display1
nary_ageclass_victimimmature:Binary..
ageclass_victimimmature:B
b_Binary.P1
b_Binary.P1
sd_Aggressor__Intercept
sd_Aggressor__Intercept
b_Binary_ageclass_victimimmature
nary_ageclass_victimimma
sd_Victim__Intercept
sd_Victim__Intercept
Chain
1
2
3
4
Chain
1
2
3
4

Model 1.3

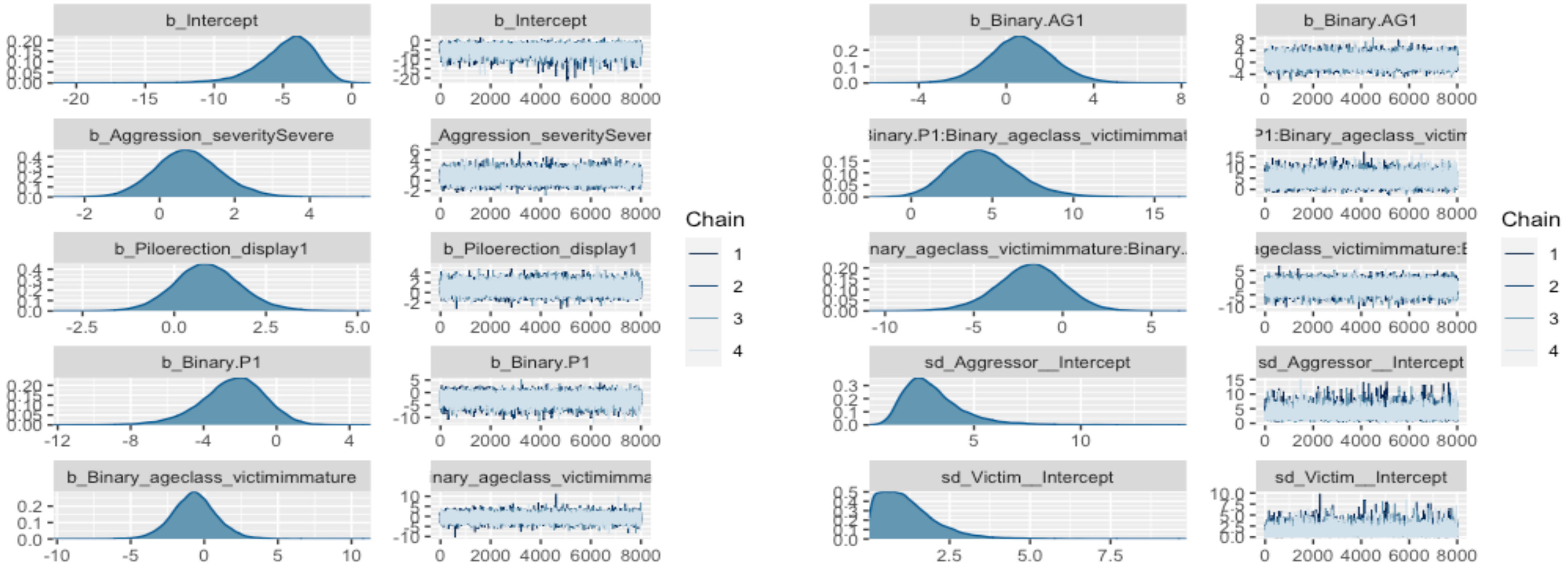
b_Intercept
b_Intercept
b_Binary.AG1
b_Binary.AG1
b_Aggression_severitySevere
_Aggression_severitySever
inary.P1:Binary_ageclass_victimimmat
P1:Binary_ageclass_victim
Chain
1
2
3
4
b_Piloerection_display1
b_Piloerection_display1
nary_ageclass_victimimmature:Binary..
ageclass_victimimmature:E
b_Binary.P1
b_Binary.P1
sd_Aggressor__Intercept
sd_Aggressor__Intercept
b_Binary_ageclass_victimimmature
nary_ageclass_victimimma
sd_Victim__Intercept
sd_Victim__Intercept

Model 1.4

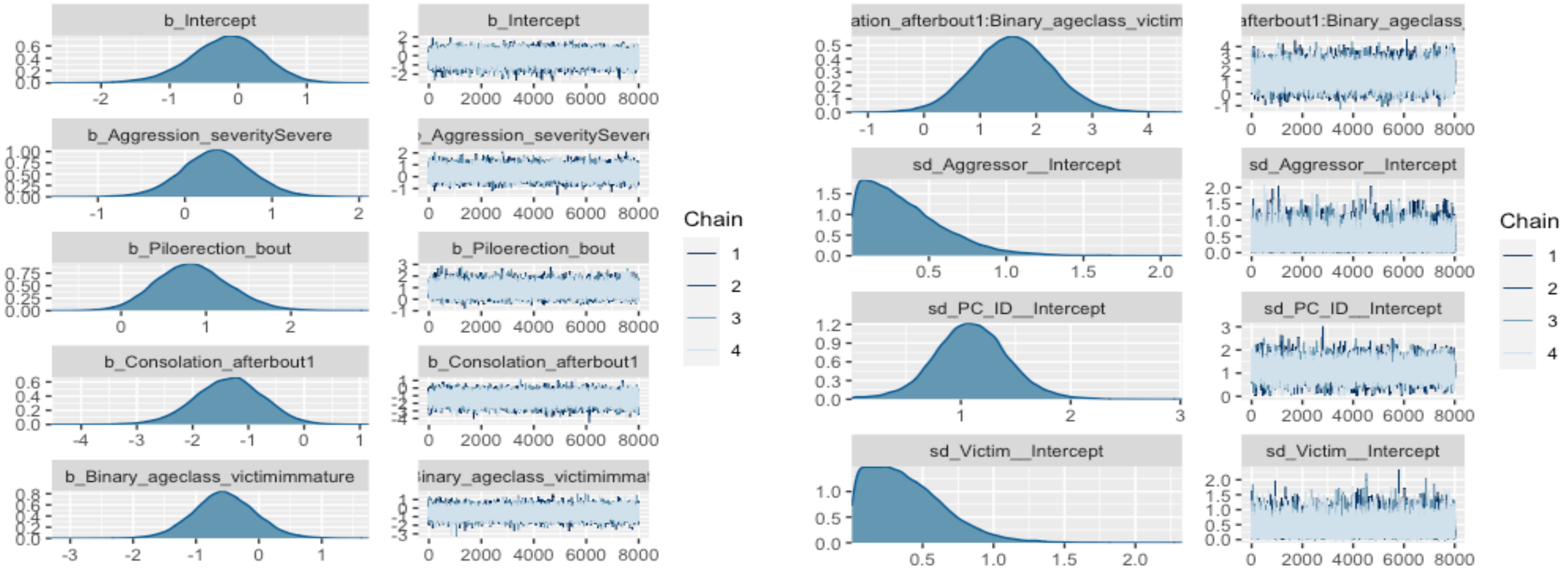
b_Intercept
b_Aggression_severitySevere
b_Piloerection_bout
b_Consolation_afterbout1
b_Binary_ageclass_victimimmature
ation_afterbout1:Binary_ageclass_victim
afterbout1:Binary_ageclass_
sd_Aggressor__Intercept
sd_PC_ID__Intercept
sd_Victim__Intercept
Chain
1
2
3
4
0 2000 4000 6000 8000

Model 1.5

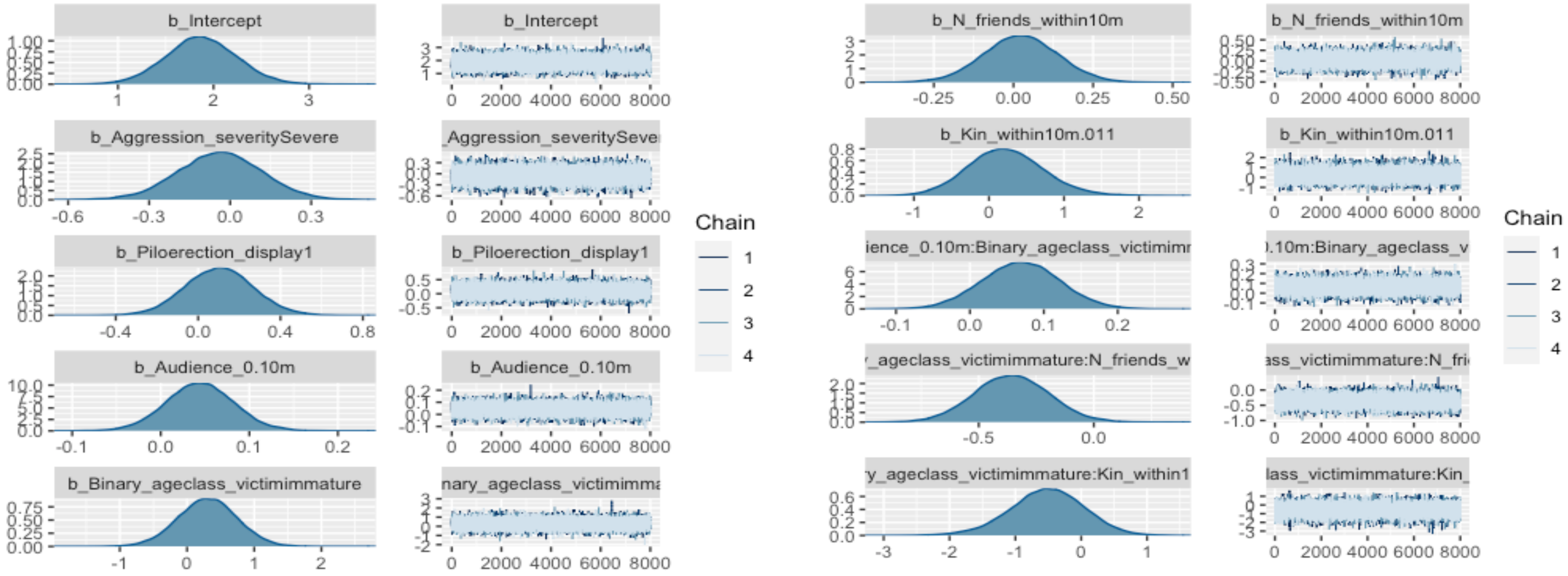

b_Intercept
b_Aggression_severitySevere
b_Piloerection_display1
b_Audience_0.10m
b_Binary_ageclass_victimimmature
b_N_friends_within10m
b_Kin_within10m.011
Chain
1
2
3
4

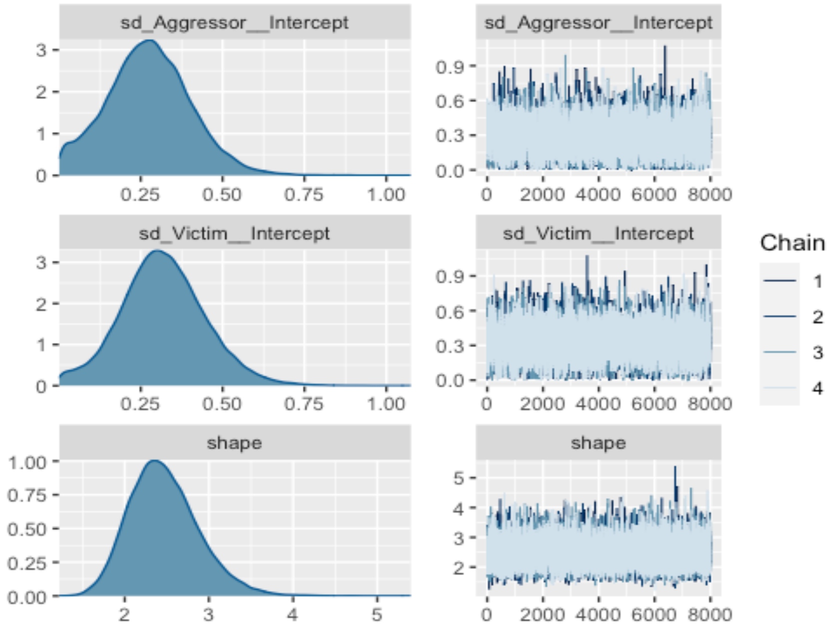
sd_Aggressor__Intercept
sd_Aggressor__Intercept
sd_Victim__Intercept
sd_Victim__Intercept
shape
shape
Chain
1
2
3
4
0.25
0.50
0.75
1.00
0
2000
4000
6000
8000

Model 1.6

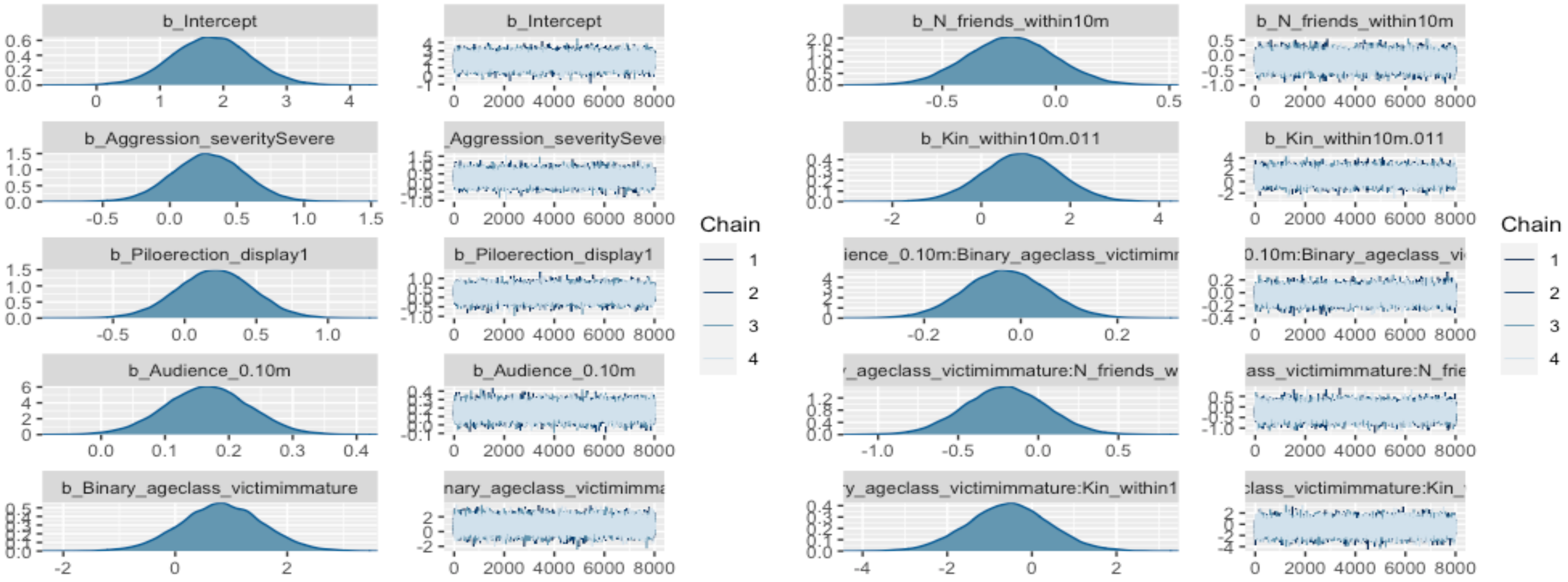

b_Intercept
b_Intercept
b_N_friends_within10m
b_N_friends_within10m
b_Aggression_severitySevere
_Aggression_severitySeve
b_Kin_within10m.011
b_Kin_within10m.011
b_Piloerection_display1
b_Piloerection_display1
ience_0.10m:Binary_ageclass_victimimr
0.10m:Binary_ageclass_vi
b_Audience_0.10m
b_Audience_0.10m
_ageclass_victimimmature:N_friends_w
ass_victimimmature:N_frie
b_Binary_ageclass_victimimmature
nary_ageclass_victimimma
y_ageclass_victimimmature:Kin_within1
class_victimimmature:Kin_
Chain
1
2
3
4
Chain
1
2
3
4

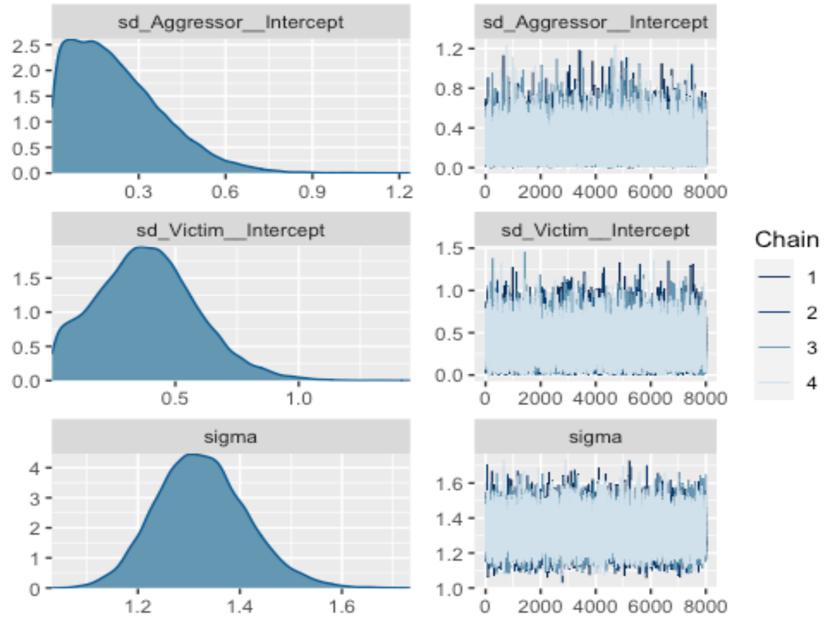
sd_Aggressor__Intercept
sd_Aggressor__Intercept
sd_Victim__Intercept
sd_Victim__Intercept
sigma
sigma
Chain
1
2
3
4

**Fig S2.** Density estimates of the observed, empirical data set *y* (black curves), with density estimates for 1000 simulated data sets $y_{rep}$ drawn from the posterior predictive distribution (bright blue curves) [100].

Model 1.1

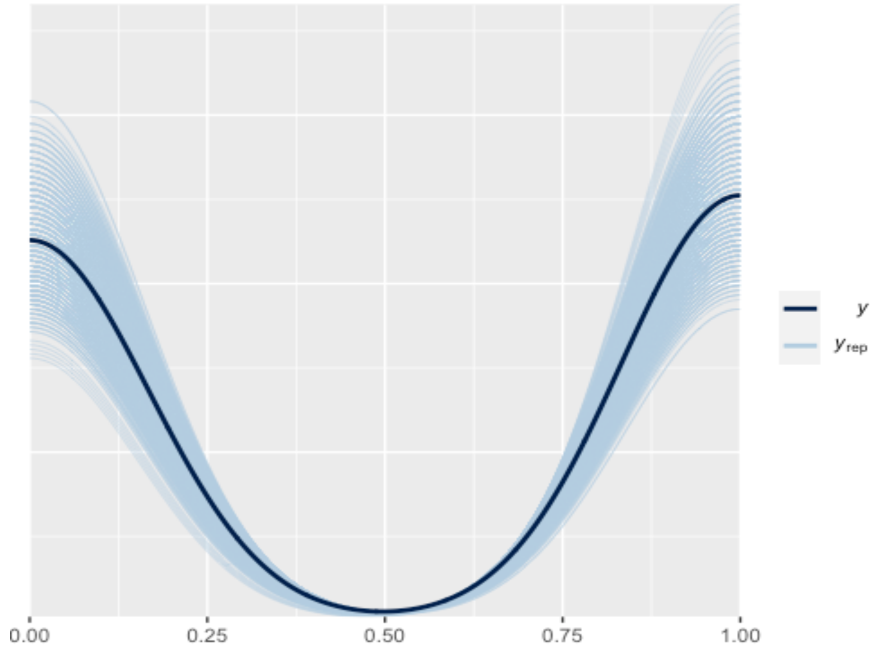

Model 1.2 **estimated response does not perfectly match empirical response due to a small sample of reconciliatory events in adults.*

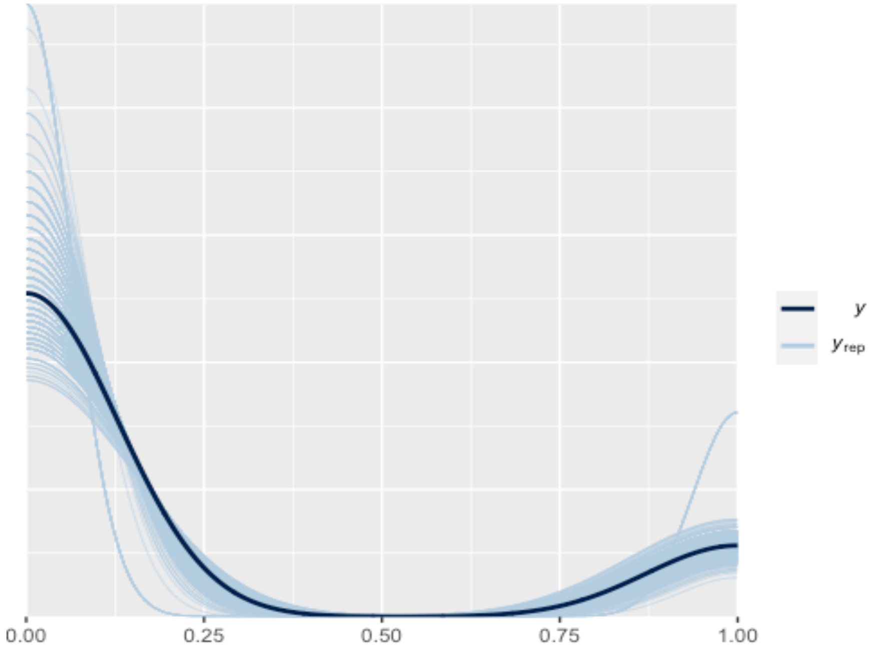

Model 1.3

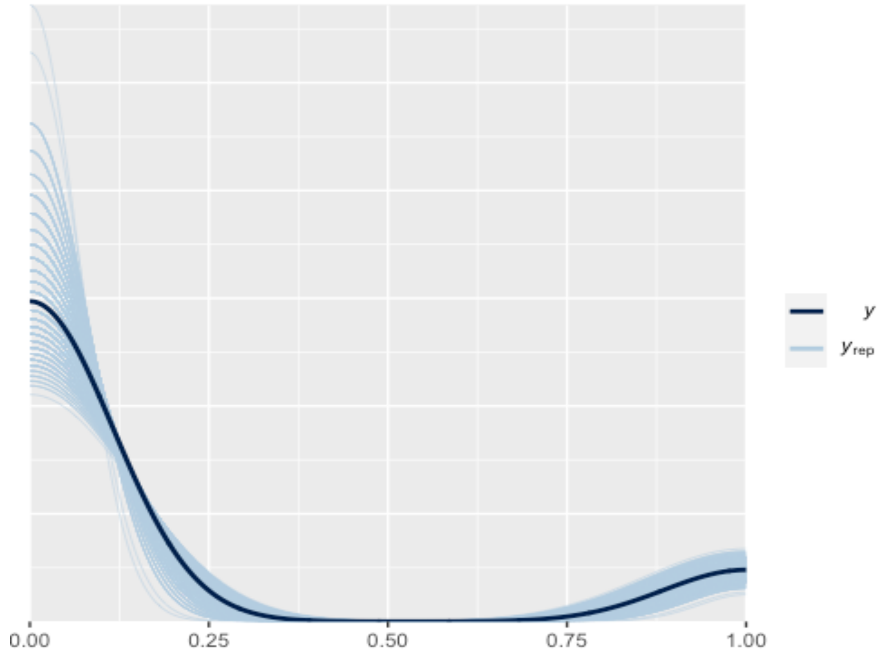

Model 1.4

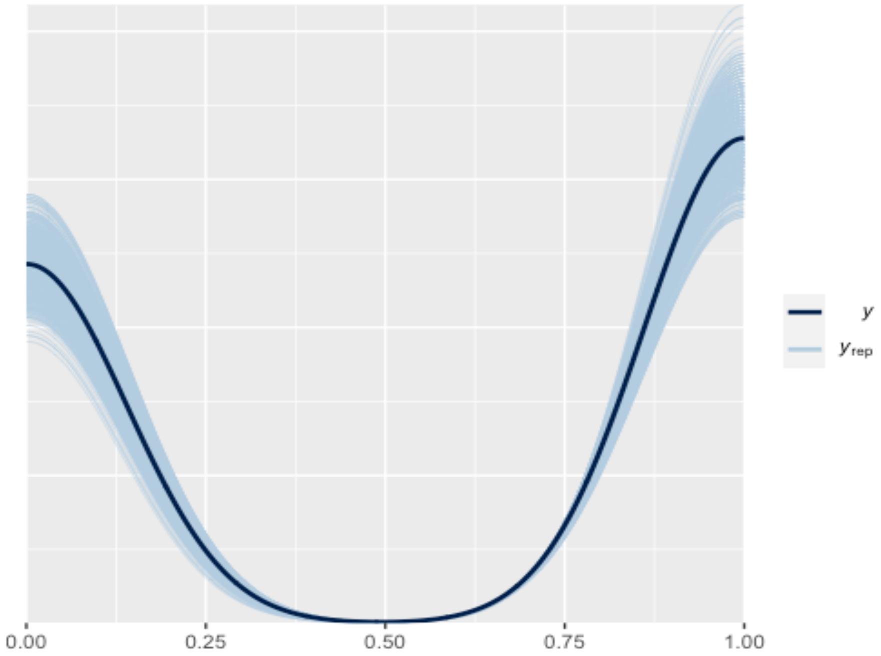

Model 1.5

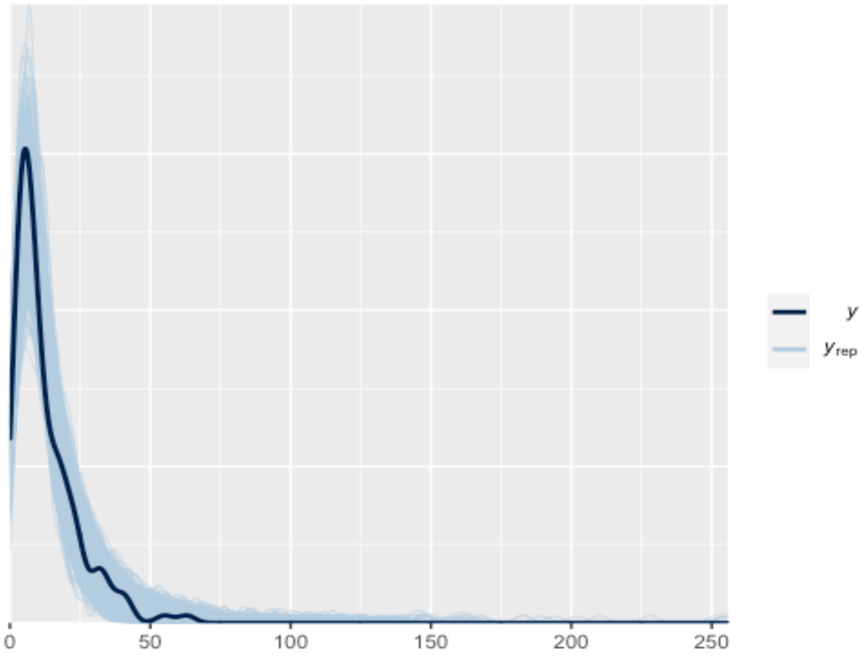

Model 1.6

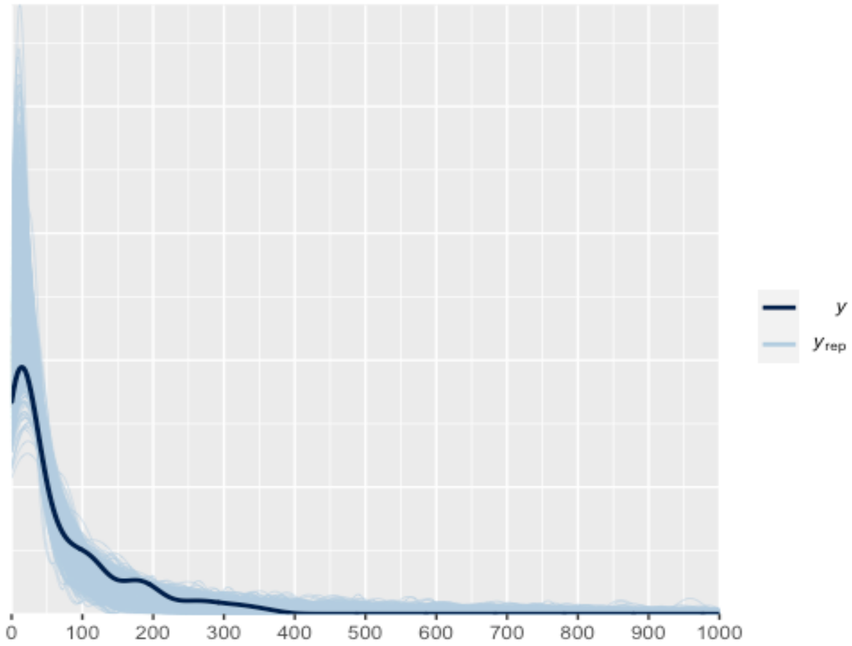

**Fig S3.** Proportion of signalling persistence after reconciliation was received or not received.

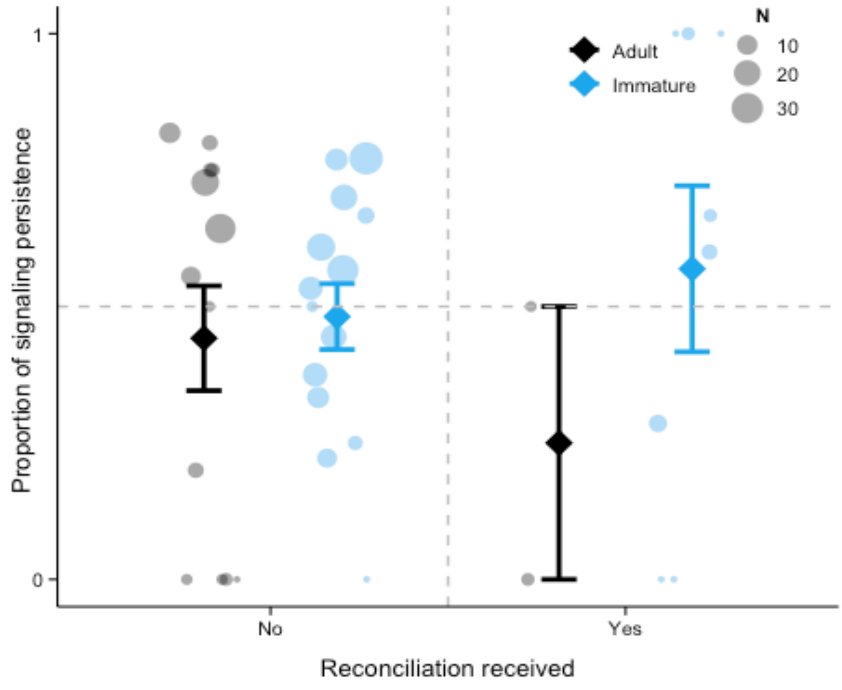

**Supplementary Image S1.** Piloerected victim.

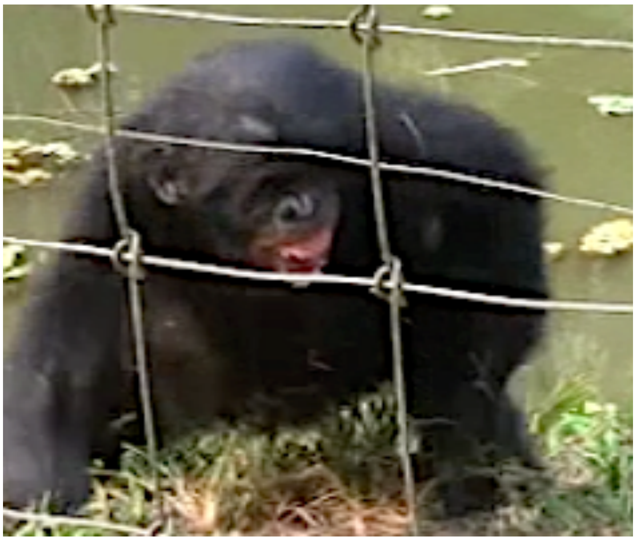

**Supplementary audio files**

Victimscream_MS.wav

Peepshriek_LM.wav

Poutmoan_YL.wav

Poutwhimper_MLK.wav

Threatbark_SL.wav

Peep_KT.wav

Contesthoot_AP.wav

Agonisticpeep_MU.wav

**Supplementary movie files**

Video descriptions

**Movie s1. Post-conflict communication in an immature bonobo victim without consolation, reconciliation or renewed aggression.**

Variables relevant to analysis:

- Original source file: PC43_EL_07.06.12.mov
- Victim ID: EL
- Victim binary age class: immature
- Victim rearing: orphaned
- Victim sex: male
- Aggressor ID: IB
- Aggression severity: mild
- Consolation received: No
- Reconciliation: no
- Renewed aggression received: no
- Number of friends within 10 m: 3
- Number of audience members within 10 m: 14
- Signal display duration: 16.3 sec
- Number of signals: 6
- Signal component types: vocalisations (2), facial expressions (4), gestures (0), body signals (0)
- Signal style categories: affiliative-submissive (6), paedomorphic (0), aggressive (0)
- Victim piloerected: No
- Out of sight events (*N*): NA
- Number bouts: 1 (no persistence)

**Movie s2. Post-conflict communication in an adult bonobo victim with consolation, but without reconciliation or renewed aggression.**

Variables relevant to analysis:

- Original source file: PC185_LS_20.07.11.mov
- Victim ID: LS
- Victim binary age class: adult
- Victim rearing: mother-reared
- Victim sex: female
- Aggressor ID: OP
- Aggression severity: severe
- Consolation received: yes
- Reconciliation: no
- Renewed aggression received: no
- Number of friends within 10 m: 4
- Number of audience members within 10 m: 19
- Signal display duration: 18.3 sec
- Number of signals: 5
- Signal component types: vocalisations (2), facial expressions (1), gestures (0), body signals (2)
- Signal style categories: affiliative-submissive (3), paedomorphic (2), aggressive (0)
- Victim piloerected: yes
- Out of sight events (*N*): 0
- Number bouts: 1 (no persistence)

**Movie s3. Post-conflict communication in an immature bonobo victim with consolation, but without reconciliation or renewed aggression.**

Variables relevant to analysis:

- Original source file: PC38_SK_23.06.11.mov
- Victim ID: SK
- Victim binary age class: immature
- Victim rearing: orphaned
- Victim sex: female
- Aggressor ID: MY
- Aggression severity: severe
- Consolation received: yes
- Reconciliation: no
- Renewed aggression received: no
- Number of friends within 10 m: 2
- Number of audience members within 10 m: 7
- Signal display duration: 14.0 sec
- Number of signals: 10
- Signal component types (*N*): vocalisations (5), facial expressions (1), gestures (1), body signals (3)

- Signal style categories (*N*): affiliative-submissive (3), paedomorphic (7), aggressive (0)
- Victim piloerected: no
- Out of sight events (*N*): 0
- Number bouts: 1 (no persistence)

**Movie s4. Post-conflict communication in an immature bonobo victim with consolation and renewed aggression, but without reconciliation.**

Variables relevant to analysis:

- Original source file: PC66_MS_25.06.11.mov
- Victim ID: MS
- Victim binary age class: immature
- Victim rearing: orphaned
- Victim sex: female
- Aggressor ID: KL
- Aggression severity: mild
- Consolation received: yes
- Reconciliation: no
- Renewed aggression received: yes
- Number of friends within 10 m: 5
- Number of audience members within 10 m: 13
- Signal display duration: 259.99 sec
- Number of signals: 24
- Signal component types (*N*): vocalisations (9), facial expressions (11), gestures (0), body signals (4)
- Signal style categories (*N*): affiliative-submissive (18), paedomorphic (6), aggressive (0)
- Victim piloerected: no
- Out of sight events (*N*): 1
- Number bouts: 6 (persisted in 3 renewed bouts after consolation was received)

**Movie s5. Post-conflict communication in an immature bonobo victim with consolation and reconciliation, but without renewed aggression.**

Variables relevant to analysis:

- Original source file: PC291_WK_06.08.11.mov
- Victim ID: WK
- Victim binary age class: immature
- Victim rearing: orphaned
- Victim sex: female
- Aggressor ID: EK
- Aggression severity: severe
- Consolation received: yes
- Reconciliation: yes
- Renewed aggression received: no

- Number of friends within 10 m: 4
- Number of audience members within 10 m: 15
- Signal display duration: 26.1 sec
- Number of signals: 8
- Signal component types (*N*): vocalisations (3), facial expressions (4), gestures (0), body signals (1)
- Signal style categories (*N*): affiliative-submissive (8), paedomorphic (0), aggressive (0)
- Victim piloerected: no
- Out of sight events (*N*): 0
- Number bouts: 3 (persisted in 2 renewed bouts after reconciliation, persisted in 0 renewed bouts after consolation was received)

**Supplementary data**

Data_and_R_script.zip (published under figshare.com, private link:

https://figshare.com/s/7dddfc02c919ec4574ef.).
